# Assessing *Zymoseptoria tritici* Candidate Effectors in the Non-Host *Nicotiana benthamiana* Reveals Immune Recognition and Potential Host-Independent Activities

**DOI:** 10.1101/2025.05.20.655180

**Authors:** Sandra V. Gomez-Gutierrrez, Catalina Rodriguez-Diaz, Namrata Jaiswal, Michael Gribskov, Matthew Helm, Stephen B. Goodwin

## Abstract

*Zymoseptoria tritici* is a significant wheat pathogen responsible for Septoria tritici blotch (STB) disease. Despite its economic impact, understanding of the molecular interactions between *Z. tritici* and its host remains limited, particularly the functions of many uncharacterized candidate effectors. Here, we assessed seven *Z. tritici* candidate effectors with elevated expression during the early biotrophic phase and transition to necrotrophy by transiently expressing them in the heterologous, non-host system *Nicotiana benthamiana*, with and without their predicted signal peptides. AlphaFold structural predictions revealed that two candidates share similarity with proteins of known function: a sterol-binding protein from *Saccharomyces cerevisiae* and a necrosis-inducing effector from the apple canker pathogen *Valsa mali*. Effector-associated phenotypes did not always correlate with expression timing, and signal peptide presence significantly influenced observed activities. Because *N. benthamiana* is not a natural host of *Z. tritici*, observed responses may reflect conserved, host-independent activities of effectors or immune recognition by *N. benthamiana,* rather than their native functions in wheat. Consistent with this distinction, two candidate effectors, Mycgr3107904 and Mycgr394290, induced cell death in *N. benthamiana* while also modulating ROS burst, likely indicative of immune recognition. Some candidates attenuate ROS production, suggesting potential targeting of conserved cellular processes, whereas none suppress PBR1-mediated cell death. Overall, our results indicate that candidate effector activities cannot be inferred from expression profiles alone, and that responses in the non-host *N. benthamiana* system reflect a combination of immune recognition and potential conserved activities, and primarily serve to prioritize candidates for validation in the native wheat–*Z. tritici* interaction.

## 1. INTRODUCTION

In plant-pathogen interactions, successful infection relies, in part, on the ability of the pathogen to modulate host defenses. To achieve this, pathogens secrete a diverse repertoire of structurally varied molecules that alter host cell structure and function, often to the advantage of the pathogen (Win et al., 2012; Seong and Krasileva, 2023; Todd et al., 2023). Among the key actors in these interactions are small, secreted proteins (SSPs) termed effectors that allow host colonization in various ways, including manipulation of the immune responses in the host (Ma, Liu and Zamany, 2019; Fabro, 2022; Todd et al., 2023; Meile et al., 2024). Effector molecules can be perceived by plant immune receptors at both the cell surface and inside cells (Dalio et al., 2018; Helm et al., 2022; Rogers et al., 2024) to induce and modulate the two-layered surveillance system of the plant immune response, which can be conceptually divided into cell-surface immunity and intracellular immunity (Bentham et al., 2020; Chang et al., 2022) .

Cell-surface immunity is the first layer of host defense and comprises a range of immune responses, including the generation and accumulation of reactive oxygen species (ROS), the deposition of callose to reinforce the plant cell wall, and the expression of defense-related genes (Bakhat et al. 2023; Chang et al. 2022; Pruitt et al. 2021; Tian et al. 2021). During plant-fungal pathogen infections, cell-surface immunity is often activated by chitin, which acts as a pathogen-associated molecular pattern (PAMP) and is recognized by host cell surface-localized pattern-recognition receptors (PRRs) (Chang et al. 2022; Tian et al. 2021). In these interactions, pathogens have developed strategies such as the secretion of effector proteins to evade chitin-triggered immunity by protecting fungal cell walls against plant chitinases, enzymes responsible for hydrolysis of chitin present in fungal cell walls that generate chitin fragments called chito oligosaccharides (COS). Effector proteins also can interfere with the recognition of COS prior to activation of cell-surface immunity, as well as disrupt signal transduction following this activation (Bakhat et al. 2023; Crumière et al. 2022; Tian et al. 2021; Volk et al. 2019).

The most well-known and extensively studied examples of cell wall protection against chitinases and disruption of chitin recognition are the Avr4 and Ecp6 effector proteins secreted by the leaf mold fungus *Fulvia fulva* (Kohler et al., 2016), and BcLysM1, a putative LysM effector secreted by the necrotrophic fungus *Botrytis cinerea* (Crumière et al., 2022). The Avr4 effector, a lectin with a chitin-binding domain, protects fungal hyphae against plant chitinases by associating specifically with chitin present in fungal cell walls. This indirectly prevents chitin hydrolysis by interfering with substrate accessibility (Van Den Burg et al. 2006; Van Esse et al. 2007). On the other hand, both Ecp6 and BcLysM1 bind with high affinity to sequester COS, competing with host receptors for chitin binding, which diminishes chitin-triggered immune responses (De Jonge et al. 2010; Sánchez-Vallet et al. 2013; Crumière et al., 2022). If cell-surface immunity is activated, some fungal effectors target key host immune components involved in signal transduction, disrupting downstream defense responses. For example, effectors can inhibit mitogen-activated protein kinase (MAPK) cascades, interfere with vesicle trafficking, or block NADP-malic enzyme activity (Jwa and Hwang, 2017; Tintor et al., 2020). In *Nicotiana benthamiana,* NIS1 from two *Colletotrichum* species suppresses ROS production triggered by flg22 and chitin, as well as the hypersensitive response (HR), by targeting immune kinases BIK1 and BAK1 (Irieda et al., 2019).

Pathogen effectors delivered into plant cells are typically detected by nucleotide-binding leucine-rich repeat receptors (NLRs) (Li, Moreno-Pérez and Coaker, 2024). However, intracellular immune activation can also arise from the perception of apoplastic effectors by PRRs, as illustrated by the interaction between the *Cladosporium fulvum* effector Avr4 and the tomato cell-surface receptor Cf-4 (Thomas et al., 1997). In addition, PAMP recognition has been shown to enhance or reinforce NLR-mediated immunity. For example, disrupting components of the PRR signaling pathway, such as deletion of the flagellin-encoding gene (*fliC*) in *Pseudomonas syringae* pv. *tabaci* 6505, significantly reduced disease resistance mediated by RAR1 and NDR1 (Zhang et al., 2010), which are accessory proteins required for the function of many NLRs in *Arabidopsis* (Tornero et al., 2002). Collectively, these findings highlight how cell-surface immunity and intracellular immunity act in concert, mutually amplifying one another to trigger a strong and coordinated immune response in plants (Yuan, Cai and Xin, 2023).

*Zymoseptoria tritici* is a fungal pathogen in the class Dothideomycetes and the causal agent of Septoria tritici blotch (STB) on wheat. It is a strictly apoplastic pathogen that secretes its effectors into the extracellular space and does not penetrate host cells (McDonald, McDonald and Solomon, 2015; Kettles et al., 2017; Petit-Houdenot, Lebrun and Scalliet, 2021). The disease involves a prolonged asymptomatic (biotrophic) phase before abruptly transitioning to a necrotrophic phase, leading to widespread plant cell death and pycnidia formation (Fagundes et al., 2020; Goodwin et al., 2011). STB is one of the most persistent and economically damaging wheat diseases, affecting both bread and durum wheat (Petit-Houdenot, Lebrun and Scalliet, 2021).

Despite the economic importance of STB and growing insights into the role of some *Z. tritici* effectors, the dissection of the molecular interactions between this pathogen and its host remain limited, particularly the functions of many uncharacterized candidate effectors. To date, relatively few effector proteins have been functionally characterized (Motteram et al., 2009; Marshall et al., 2011; Ben M’Barek et al., 2015; Gohari, 2015; Rudd et al., 2015; Sánchez-Vallet et al., 2020; Karki et al., 2021; Tian et al., 2021). For example, three LysM domain-containing proteins (LysM) have been identified from *Z. tritici* and one, Mg3LysM, was the first to be confirmed as capable of suppressing chitin-induced defense responses in wheat and protecting fungal hyphae against chitinase hydrolysis (Marshall et al., 2011). This effector is homologous to Ecp6 from *Fulvia fulva* (Marshall et al., 2011). Mg1LysM, a functional homolog of the chitin-binding effector Avr4 from *F. fulva*, was later found to shield fungal hyphae from host chitinase-mediated hydrolysis (Sánchez-Vallet et al., 2020). A separate study demonstrated that Mgx1LysM, the third LysM effector, can bind chitin, suppress chitin-induced ROS production, and protect fungal hyphae from chitinase-mediated degradation (Tian et al., 2021). Notably, all three LysM effectors exhibit partial functional redundancy (Tian et al., 2021). Furthermore, Karki et al. (2021) recently identified *Z. tritici* effector ZtSSP2, which is expressed at 2 days after inoculation (DAI) and interacts with a wheat E3 ubiquitin ligase (TaE3UBQ). Silencing *TaE3UBQ* via virus-induced gene silencing led to increased susceptibility to *Z. tritici*, indicating that *TaE3UBQ* may play a role in regulating defense responses in wheat. This finding suggests that ZtSSP2 could suppress TaE3UBQ ligase activity, thereby enhancing *Z. tritici* pathogenicity (Karki et al., 2021).

The necrotrophic effectors ZtNIP1 and ZtNIP2 have been found to induce cell death and chlorosis in some wheat cultivars. However, their mechanism of action remains unclear (Ben M’Barek et al., 2015). ZtNIP1 contains an Hce2 domain (for **H**omologs of ***C****ladosporium fulvum* **E**cp2 effector), a conserved domain that was shown to induce plant cell death and is widely conserved in the fungal kingdom (Zhang et al., 2019). A more recent study found that some effectors in *Z. tritici* that contain an Hce2 domain are structurally related to killer proteins from *Ustilago maydis*, and the analysis of their biological activity revealed that ZtNIP1 (renamed as Zt-KP4 in Guillen et al., 2024), has a toxic activity against other fungi such as *Botrytis cinerea* and even *Z. tritici* itself (Guillen et al., 2024).

Recent analyses have employed heterologous systems, such as *Agrobacterium*-mediated expression in *N. benthamiana*, to gain insight into the potential roles and functional characterization of *Z. tritici* effectors. One study demonstrated that recognition of *Z. tritici* candidate effectors is common in *N. benthamiana* and occurs at the plasma membrane-apoplast interface, involving defense-associated receptor-like kinases (RLKs) NbBAK1 and NbSOBIR1 (Kettles et al., 2017). Screening of *Z. tritici* effector libraries in *N. benthamiana* identified those that suppress conserved immune responses, highlighting this as a useful tool for uncovering effector functions that could later be validated in wheat (Thynne et al., 2024). Although *N. benthamiana* is particularly suited for rapid functional screening of candidate effectors, it is a non host for *Z. tritici* and its immune receptor repertoire differs substantially from that of wheat. Responses observed in this system should therefore be interpreted as heterologous recognition events, which may or may not reflect the true functional activities of these effectors during infection of their natural host.

To investigate the potential roles of *Z. tritici* candidate effectors in modulating immunity responses, we selected seven genes from an RNAseq dataset of *Z. tritici* during infection of a susceptible wheat cultivar (Gomez-Gutierrez et al., 2025). These candidates were prioritized because they exhibited elevated expression during the early biotrophic phase and the transition to necrotrophy, suggesting a potential involvement in distinct stages of disease progression. Additionally, AlphaFold structural predictions revealed that two candidates share similarity with proteins of known function, providing further insights into their potential biological activity.

Here, we tested seven candidate effectors using transient expression in *N. benthamiana* both with and without their predicted signal peptides to assess whether the presence of this region influences ROS suppression activity. The constructs in our study lacking the signal peptide represent the putative mature form of the protein after signal peptide cleavage, enabling us to investigate potential intracellular activities. In contrast, the constructs retaining the signal peptide allowed us to evaluate whether the presence of this region contributes to or is required for the observed phenotype. For the assays on suppression of NLR-mediated cell death and cell death induction, we restricted our analysis to the candidate effectors expressed with their putative signal peptides, based on the biology of *Z. tritici* as an apoplastic pathogen that does not penetrate host cells, and the observation that the majority of characterized *Z. tritici* effectors function in the extracellular space across infection phases. We therefore prioritized testing the version of the candidate effectors that most closely reflects their native mode of deployment during host colonization in the follow-up assays.

We hypothesized that the activity of the seven candidate effectors, such as the suppression of chitin-triggered ROS bursts, interference with NLR-triggered cell death, or direct induction of host cell death, would reflect their stage-specific expression, and that the presence of a signal peptide would influence their function. Our results show that the activity of some candidate effectors did not align with their expression profiles at specific stages of the disease. For example, Mycgr3107904 attenuates chitin-induced ROS despite being highly expressed during the transition to necrotrophy and induces necrosis in *N. benthamiana*. Although the phenotype observed for Mycgr3107904 in *N. benthamiana* does not correspond to its expression pattern, it is consistent with the presence of its predicted Hce2 domain and its structural similarity to the *Valsa mali* effector VmHEP1, both of which are associated with necrosis induction. None of the tested effectors suppress PBR1-mediated cell death, indicating they do not target this NLR or its downstream signaling. Importantly, we confirm that the presence of a signal peptide influences the activity of candidate effectors on the suppression of chitin-triggered ROS bursts. These results provide a foundation for further research on candidate effectors from conserved protein families, which are present among several fungal species, and might have a potential role in *Z. tritici* virulence on wheat. In addition, we identify putative functions for a species-specific effector, suggesting the pathogen also may rely on unique, uncharacterized effectors to cause disease in wheat.

## 2. RESULTS

### 2.1. Conservation of candidate effectors

Given that the predicted effector repertoire in *Zymoseptoria* varies considerably both among *Z. tritici* isolates and in comparison to its sister species (Feurtey et al., 2020), we aimed to assess the conservation of 32 candidate effectors across these genomes. We obtained genome assemblies and predicted proteomes from 18 *Z. tritici* isolates and three closely related *Zymoseptoria* species (*Z. ardabiliae, Z. brevis,* and *Z. pseudotritici*) (Badet et al., 2020; Feurtey et al., 2020), and the nucleotide sequences of 32 candidate effectors from the Dutch isolate IPO323. Using tBLASTx, we found that most candidate effectors are present across all *Z. tritici* isolates and sister species (Figure 1). However, candidate effector Mycgr3106502 is abscent in the Ukranian isolate UR95, four effectors that are located on chromosome one are absent in the European isolate 3D1; candidate effector Mycgr3102792 is absent in two of the *Zymoseptoria* sister species; Mycgr383064 is missing in *Z. brevis*; and Mycgr3101652 is absent in all three *Zymoseptoria* sister species (Figure 1).

**FIGURE 1.**
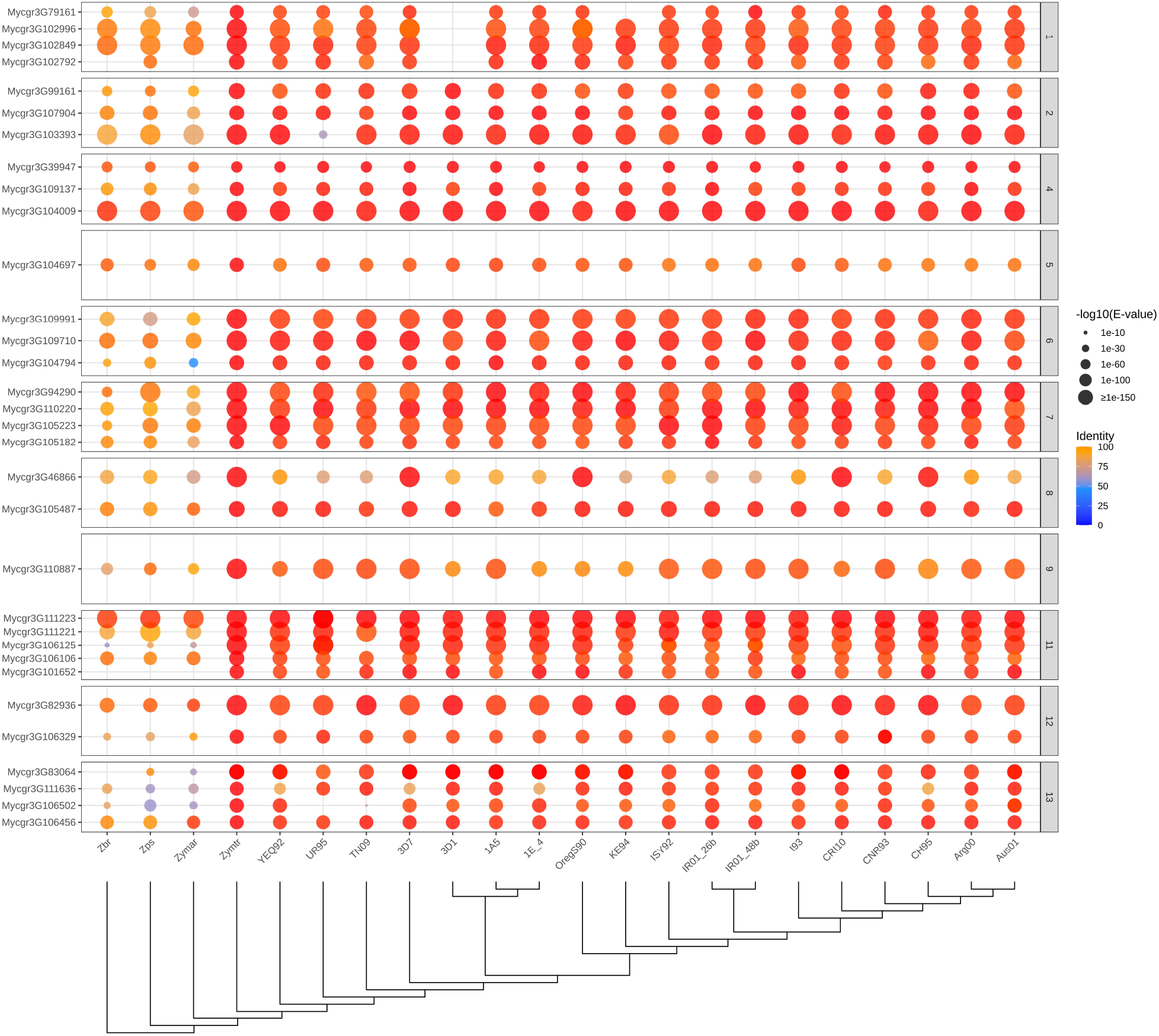
Sequence conservation of orthologs for 32 candidate effector genes across 19 isolates of *Zymoseptoria tritici* and the three sister species *Z. pseudotritici* (Zps)*, Z. brevis* (Zbr), and *Z. ardabiliae* (Zymar). Conservation was determined by tBLASTx searches against a nucleotide database containing the genome scaffolds of all 22 isolates including IPO323. The reference strain IPO323 is shown as Zymtr. Isolates are indicated at the bottom and candidate effector genes at the left side. A tree showing the phylogenetic relationships among the isolates based on all genes generated by OrthoFinder is indicated on the bottom side. Chromosome number in which each candidate effector is located is indicated on the right side as individual panels. Sequence conservation is presented as percentage of identity and it is represented as a color scale with blue showing lower percentage of identity and red showing the highest percentage of identity compared to the candidate effector sequence in the reference strain IPO323.

Notably, tBLASTx searches revealed that the majority of candidate effectors share greater than 90% sequence identity across the 18 *Z. tritici* isolates (Figure 1). In contrast, sequence variation among all 32 candidate effectors is markedly higher in the three *Zymoseptoria* sister species, where sequence identity drops to approximately 75% for most candidates. This divergence is even more pronounced for effectors encoded on chromosomes 6, 11, 12 and 13, where sequence identity falls below 60%, as observed for Mycgr3111636, Mycgr3106502, Mycgr3106125, and Mycgr3104794 (Figure 1). Remarkably, tBLASTx searches revealed that the DNA sequence of candidate effector Mycgr394290 from the Dutch isolate IPO323 is present in the corresponding genomic region of chromosome 7 across all 21 *Zymoseptoria* isolates (Figure 1). However, a BLASTp search revealed that a ∼60-amino acid region of candidate effector Mycgr394290 showed a complete match to a short, predicted protein in the European isolates 3D7, 1A5, and 1E4, but was absent from the called proteomes of all other isolates. While nucleotide-level similarity across the locus was detected using tBLASTx (Figure 1), translation of the corresponding genomic region did not yield a continuous open reading frame. Further analysis with BLASTx recovered only partial similarity (∼56% to 64% coverage) in translated nucleotide queries from sequences of isolates 3D7, 1A5, and 1E4 to the reference protein. These results indicate that the locus is likely disrupted and does not encode a full-length functional effector;, the annotated short protein likely representing a truncated remnant of the gene.

BLASTp analysis showed that the candidate effector proteins Mycgr3109710 and Mycgr3103393 each matched two distinct proteins across the 21 isolates. One match showed >98% sequence identity, while the other showed 53–58% identity. Both candidates are annotated as Pathogenesis-Related 1 (PR1) proteins, and *Z. tritici* strains typically encode three to four PR1 proteins. Mycgr3109710 and Mycgr3103393 are more similar to each other than to the other PR1 proteins encoded in *Z. tritici* (data not shown). Therefore, the two BLASTp matches likely correspond to these two proteins: each candidate matches its ortholog in all isolates (>98% identity) as well as the other PR1 protein (∼54% identity). We observed that while the predicted protein sequence of Mycgr3109710 is fully conserved across all 21 *Zymoseptoria* isolates (98% to 100% protein sequence identity), the protein sequence of Mycgr3103393 was absent in isolate UR95 but shared 53% identity with the orthologue of Mycgr3109710 in isolate UR95, which is also supported by the tBLASTx results that show less than 60% sequence identity for Mycgr3103393 in the Ukrainian isolate UR95.

### 2.2. AlphaFold predicted structures of candidate effectors from *Z. tritici*

Given that only 13 of the 32 selected candidate effectors have functional annotations, and considering that effectors often lack sequence similarity due to their rapid divergence and host specialization (Sperschneider et al., 2015), predicting the tertiary structures of fungal effectors is a valuable approach for gaining insights into their potential functional roles and evolutionary trajectories (Seong and Krasileva, 2023). To investigate the potential functions of candidate effectors, we predicted their tertiary structures using AlphaFold3 (Abramson et al., 2024) and analyzed these structures for similarities to proteins with known tertiary structures and, in some cases, characterized functions, using both the FoldSeek server (https://search.foldseek.com/search) (Van Kempen et al., 2024) and the DALI server (Holm, 2022). Predicted Aligned Error (PAE) plots were generated (Supplementary figure 1A) to assess prediction reliability. Effectors were grouped in Figure 2 based on a multiple structural alignment using FoldMason MSA (https://search.foldseek.com/foldmason) (Supplementary figure 1B), with each row representing a cluster in the tree generated by the MSA.

**FIGURE 2.**
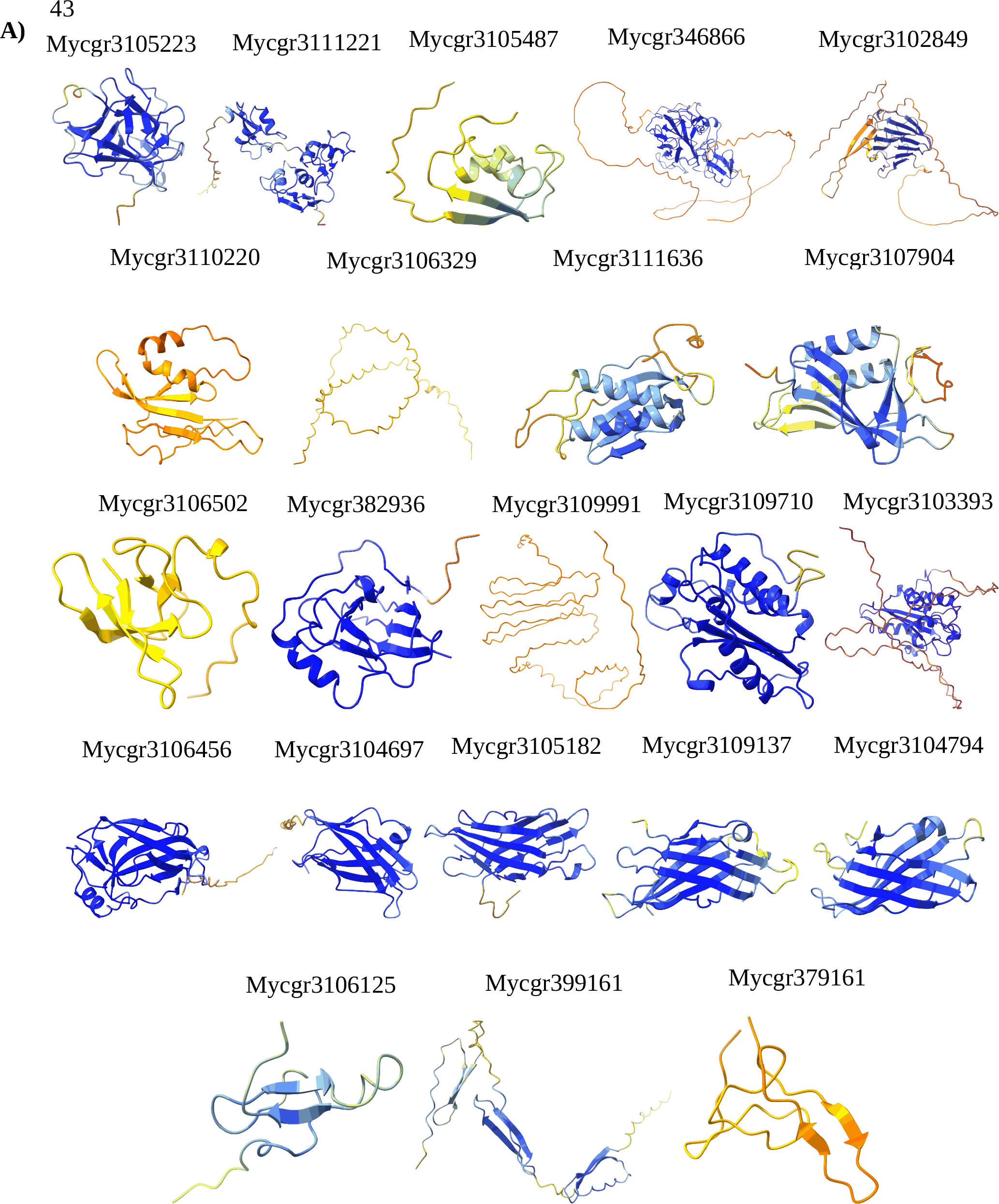
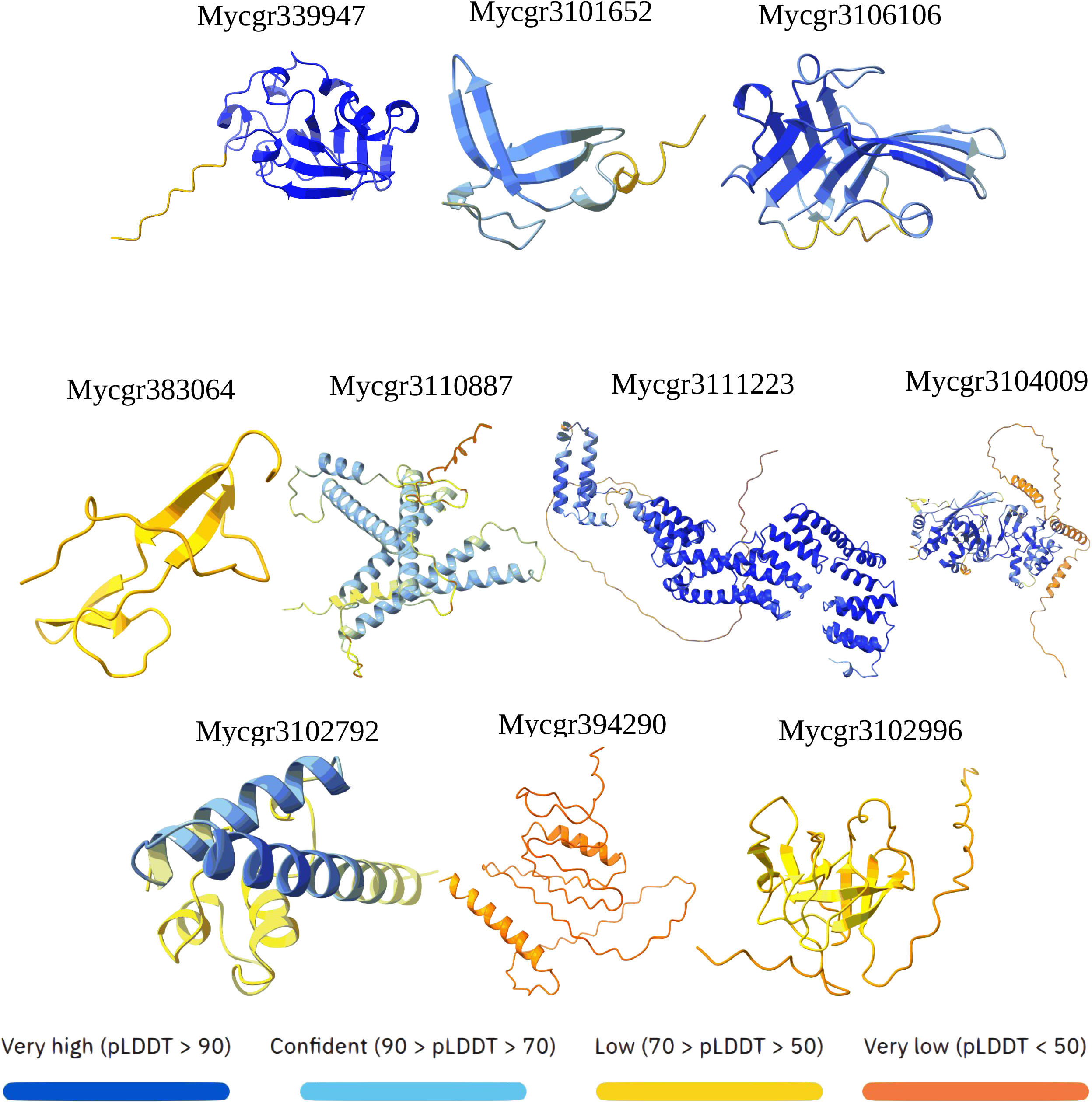
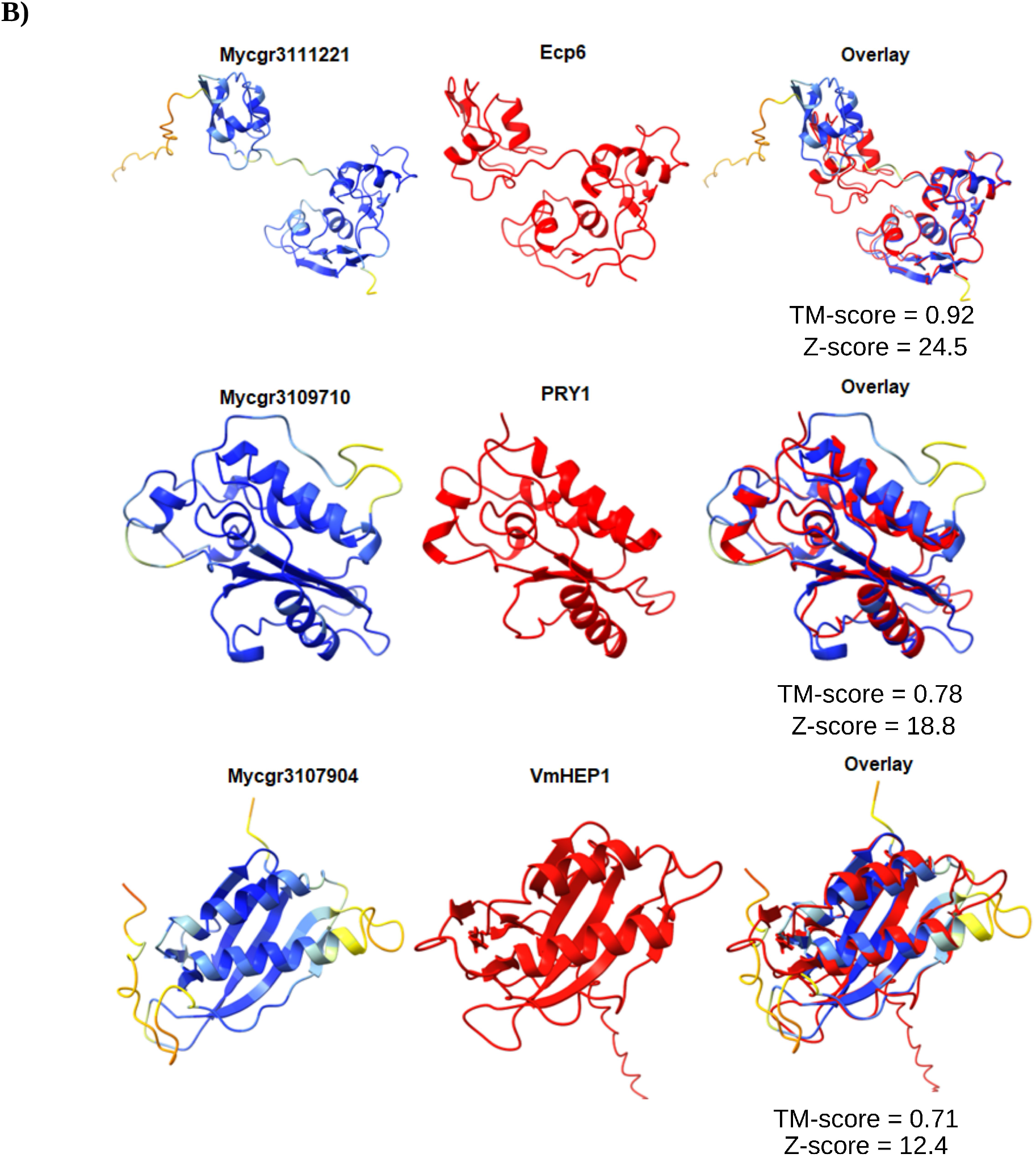
Predicted structures of 32 selected candidate effector proteins in the wheat pathogen *Zymoseptoria tritici* using AlphaFold v3.0. A) Effectors were grouped based on a multiple structural alignment using FoldMason MSA (https://search.foldseek.com/foldmason). Each row represents a cluster within the tree generated by the MSA. B) Alignments of three candidate effectors with structural similarity to characterized proteins from *Fulvia fulva, Saccharomyces cerevisiae* and *Valsa mali* as determined by FoldSeek analysis. The *Z. tritici* effector is on the left, the corresponding effector being compared is in the middle, and the overlay of both effectors is on the right. TM-scores from Fold-Seek and Z-scores from DALI were included for each structural alignment.

Structural modeling predictions revealed that most candidate effectors contain regions with either very high confidence (pLDDT > 90) or high confidence (pLDDT 70–90) (Figure 2A, Supplementary figure 1A). Notably, the amino- and carboxy-terminal regions in nearly all effectors were predicted with very low confidence (pLDDT < 50) (Figure 2A, Supplementary figure 1A).

The high confidence in the structural prediction of specific domains within some candidate effectors in our set, which possess functional annotations and experimentally validated roles, highlights the reliability of tertiary structure modeling. For example, the LysM domains predicted within the candidate effector Mycgr3111221 were modeled with very high confidence (pLDDT > 90) (Figure 2A). Structural similarity analysis using FoldSeek showed that Mycgr3111221 closely resembles the *Fulvia fulva* (formerly *Cladosporium fulvum*) LysM effector Ecp6, with an E value of 3.13 × 10^-28^ and a TM score of 0.92. Consistently, the DALI structural alignment produced a Z score of 24.5 (Supplementary Table 2). Ecp6 has previously been reported as a close homolog of Mycgr3111221 (also known as Mg3LysM) (Marshall et al., 2011). Both effectors contain three LysM domains, with two of these domains located toward the C-terminal region of the predicted protein (Marshall et al., 2011). These proteins share 48.6% amino acid sequence identity. Similarly, candidate effector Mycgr3105487 (also known as Mg1LysM), which contains a single LysM domain and has been experimentally shown to protect cell wall chitin from hydrolysis by plant enzymes (Marshall et al., 2011; Sánchez-Vallet et al., 2020), exhibited significant structural similarity to the *F. fulva* effector Ecp6, with an E value of 1.46 × 10^-4^ and a TM score of 0.78, particularly in the folding pattern of the single LysM domain in Mycgr3105487. Notably, Mycgr3105487 shares 41.7% amino acid sequence identity with Ecp6 as obtained from the FoldSeek output. Consistently, the DALI structural alignment produced a Z score of 8.6 (Supplementary Table 2).

Structural similarity analyses indicated that the candidate effector Mycgr3109710 is likely to adopt a protein fold closely resembling the CAP-domain–containing protein PRY1 from *Saccharomyces cerevisiae* (Figure 2B). This inference is supported by both FoldSeek structural alignment results, which yielded an E value of 1.81 × 10^-11^ and a TM score of 0.78, and DALI structural alignment that produced a Z score of 18.8 (Figure 2B). Additionally, significant structural similarities were identified with multiple PR1-L proteins from other fungal pathogen species, including *F. fulva*, *Cercospora zeae-maydis*, *Venturia nashicola*, and *Pyricularia oryzae*. Many of these proteins are annotated as putative effectors. This finding highlights the structural conservation of Mycgr3109710 as a PR1-L protein among PR-1 family members, whose functions remain unknown, across fungal species with varying lifestyles within the same taxonomic class.

The candidate effector Mycgr3107904 was predicted to adopt a protein structure similar to that of putative effectors containing an Hce2 (**H**omologs of ***C****. fulvum* **E**cp2 effector) domain (Figure 2B), found in fungal plant pathogens such as the apple pathogen *Colletotrichum noveboracense* (E-value: 7.81 × 10^-8^), *Aspergillus* sp.(multiple matches), *Fusarium oxysporum* (E value: 2.41 × 10^-8^) and the apple canker pathogen *Valsa mali* (E value: 2.99 × 10^-5^). Structural similarity searches using FoldSeek further confirmed that Mycgr3107904 exhibits conserved folding patterns among other *Z. tritici* isolates, including the European isolate 1E4 (E value: 2.88 × 10^-^12_)._

To explore potential functional relevance, we investigated a specific Hce2 domain-containing effector named VmHEP1, which has been characterized extensively and contributes to the virulence of the apple pathogen *V. mali* (Zhang et al., 2019). We used AlphaFold v3.0 to predict the tertiary structure of VmHEP1 and performed pairwise structural alignments against the predicted structure of Mycgr3107904 using the RCSB PDB tool (https://www.rcsb.org/alignment) (Zardecki et al., 2016). Despite only 17% sequence identity, we found that VmHEP1 is a putative structural homolog of Mycgr3107904 (Figure 2B), with a TM score of 0.71 obtained from the RCSB PDB tool. We also used the DALI pairwise structure comparison tool (Holm, 2022) and obtained a Z score of 12.4 for the structural alignment between VmHEP1 and Mycgr3107904. This is remarkable, as studies have highlighted the role of Hce2 domain-containing effectors as necrosis-inducing proteins, or in suppression of cell surface-triggered immunity responses (Laugé et al., 1997; Ben M’Barek et al., 2015; Thynne et al., 2024). Additionally, proteins with the Hce2 domain in certain fungal species, including *Z. tritici*, are associated with a potential role in inhibiting fungal competitors (Stergiopoulos et al., 2012; Guillen et al., 2024). This potential dual functionality underscores the diverse roles of Hce2-domain-containing proteins in fungal biology and pathogenicity.

### 2.3. Some candidate effectors of *Z. tritici* modulate chitin-mediated ROS production depending on the presence of their signal peptides

To gain initial insight into the potential functions of the selected effector candidates, we investigated whether they modulate the production of reactive oxygen species (ROS) by transiently expressing them in *N. benthamiana*, both with and without their predicted signal peptides.

Contrary to our hypothesis, the transient expression of two candidate effectors with early gene expression, Mycgr3109710 and Mycgr3111221 (without their signal peptides), increased ROS production compared to the control (free sYFP + chitin), rather than attenuating the maximum chitin-mediated ROS production across three distinct biological replicates (Figure 3A). However, according to the statistical analyses, their effect was not significant compared to the control free sYFP. In contrast, the transient expression of two candidate effectors, Mycgr394290 and Mycgr3107904 (without their signal peptides), consistently attenuated chitin-mediated ROS production compared to the control in *N. benthamiana* (Figure 3A) and their effect was significant (p value < 0.05). Furthermore, three candidate effectors, Mycgr3104794, Mycgr3103393, and Mycgr3106502, showed no effect on ROS production when expressed transiently in *N. benthamiana* cells without their signal peptides, as their activity neither increased nor attenuated ROS levels relative to the control (Figure 3A).

**FIGURE 3.**
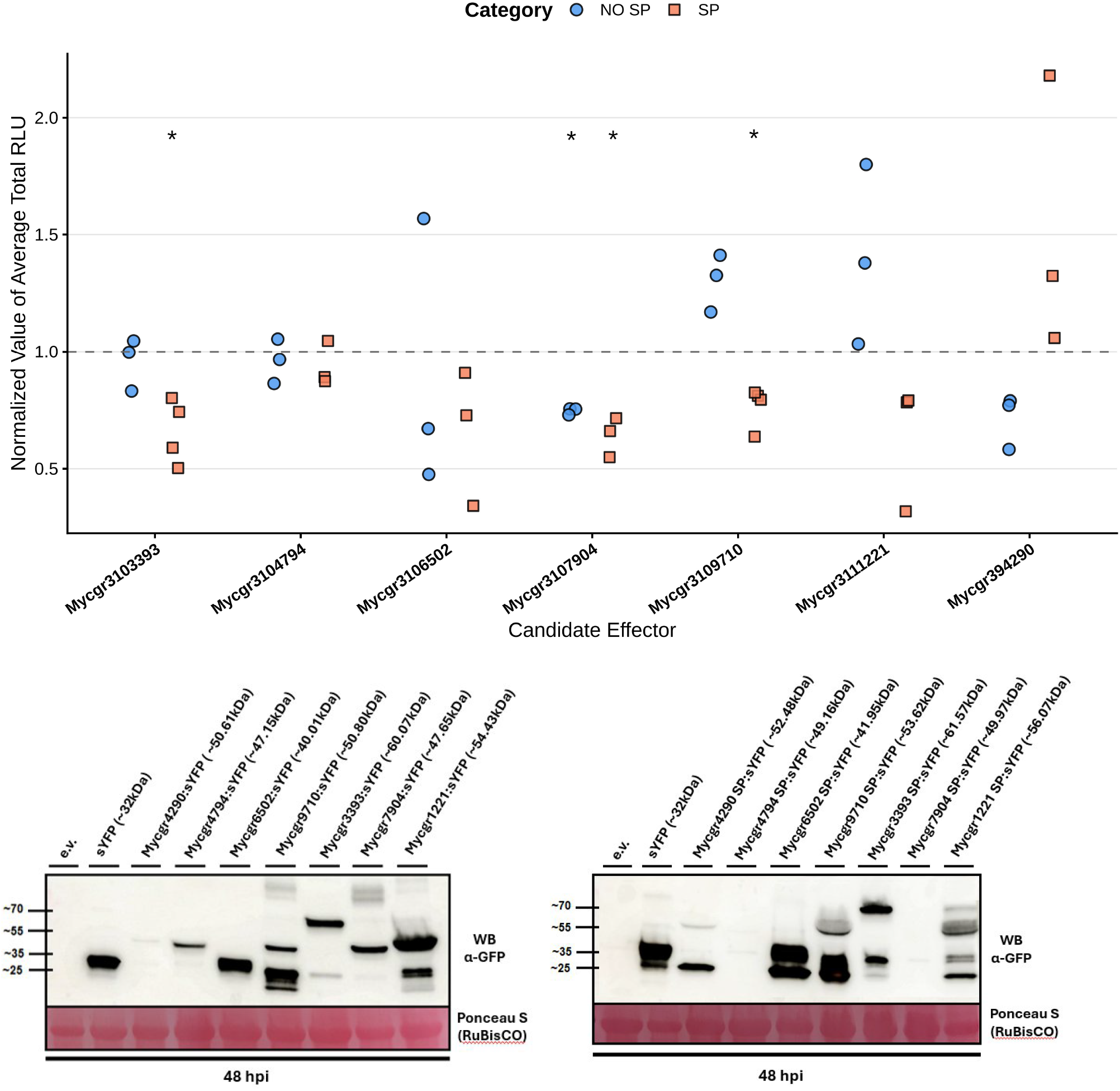
Production of Reactive Oxygen Species (ROS) and protein accumulation of *Zymoseptoria tritici* candidate effectors in *Nicotiana benthamiana*. A) Production of ROS in *N. benthamiana* leaves infiltrated with *Agrobacterium tumefaciens* strain GV3101 carrying the *Z.* tritici candidate effector genes without the signal peptide (blue) and with their native signal peptides (orange). For each biological replicate, the relative luminescence accumulation (RLU) was measured for 12 individual leaf disks over 40 minutes in a microplate reader. The average total RLU from all of the leaf discs in each ROS assay was calculated and normalized by dividing the average total luminescence of effector-expressing leaf discs to the average of the leaf discs expressing the negative control free sYFP fluorescent protein. Three to four biological replicates were included for each candidate effector. B) Immunoblot analyses of the *Z. tritici* effector-fluorescent protein fusions expressed without their signal peptides (left) and with their signal peptides (right) showing protein accumulation.

In contrast to the results observed when expressing Mycgr3109710 without its native signal peptide, transient expression of this effector with its signal peptide consistently attenuated chitin-mediated ROS production compared to the free sYFP control across four independent biological replicates (Figure 3A). Similarly, transient expression of effector Mycgr3111221 including its putative signal peptide consistently attenuated ROS production across three biological replicates but was not statistically significant when compared to the control free sYFP. However, the trend of attenuation in ROS production coincides with previous reports on the activity of this effector (Tian et al., 2021).

Notably, candidate effector Mycgr3107904 attenuates ROS production and showed a significant effect (p-value < 0.05) compared to the control free sYFP when expressed with its putative signal peptide, as it did when expressed without its signal peptide (Figure 3A). This suggests that the activity of this effector in modulating plant immune responses may be independent of its localization.

Transient expression of Mycgr3103393 with its signal peptide significantly attenuated ROS production (Figure 3A), in contrast to its lack of effect when the signal peptide was removed. Contrary to what we observed when expressing Mycgr394290 without its signal peptide, this candidate effector (with its native signal peptide) increased ROS production compared to the free sYFP control but the effect was not statistically significant due to the high variability across the biological replicates. Additionally, one candidate effector, Mycgr3104794, showed no significant effect on ROS production when expressed transiently in *N. benthamiana* cells with its signal peptide, as its activity neither increased nor attenuated ROS levels relative to the control (Figure 3A). We observed protein accumulation for all seven tested effectors, with Mycgr394290 expressed without its signal peptide, and Mycgr3104794 and Mycgr3107904 expressed with their native signal peptides showing a light band that indicates lower protein accumulation compared to the control free sYFP and the rest of the candidate effectors. However, all the effectors both expressed with and without their signal peptides showed the expected band size in the western blots (Figure 3B).

### 2.4. None of the tested *Z. tritici* effectors interfere with PBR1-D496V-induced cell death

Given the observed ROS attenuation activity of some candidate effectors when transiently expressed with their signal peptides, we hypothesized that these proteins might also suppress NLR-mediated cell death signaling. To test this possibility, we used the DEX-inducible autoactive PBR1(D496V) system as a robust and experimentally tractable assay for NLR-dependent hypersensitive responses. With this approach, we expected to assess whether any of the tested candidate effectors interfere with NLR-triggered cell death pathways.

Contrary to our expectations, none of the tested candidate effectors consistently suppressed PBR1(D496V)-induced cell death one day after DEX application (three days post infiltration), as shown in Figure 4A. Mycgr3111221 was excluded from this analysis because its primary function has been associated previously with protection of fungal hyphae against plant chitinases rather than cell death suppression. The results were consistent across three independent replicates, suggesting that none of the selected effectors, when expressed with their putative signal peptides, directly targets PBR1 or any of its downstream signaling components after activation. However, it remains possible that these effectors act by interfering with effector recognition or interactions with helper proteins. These upstream events were not tested in our experimental setup, as the autoactive PBR1(D496V) variant used here is constitutively active and does not require AvrPphB for activation.

**FIGURE 4.**
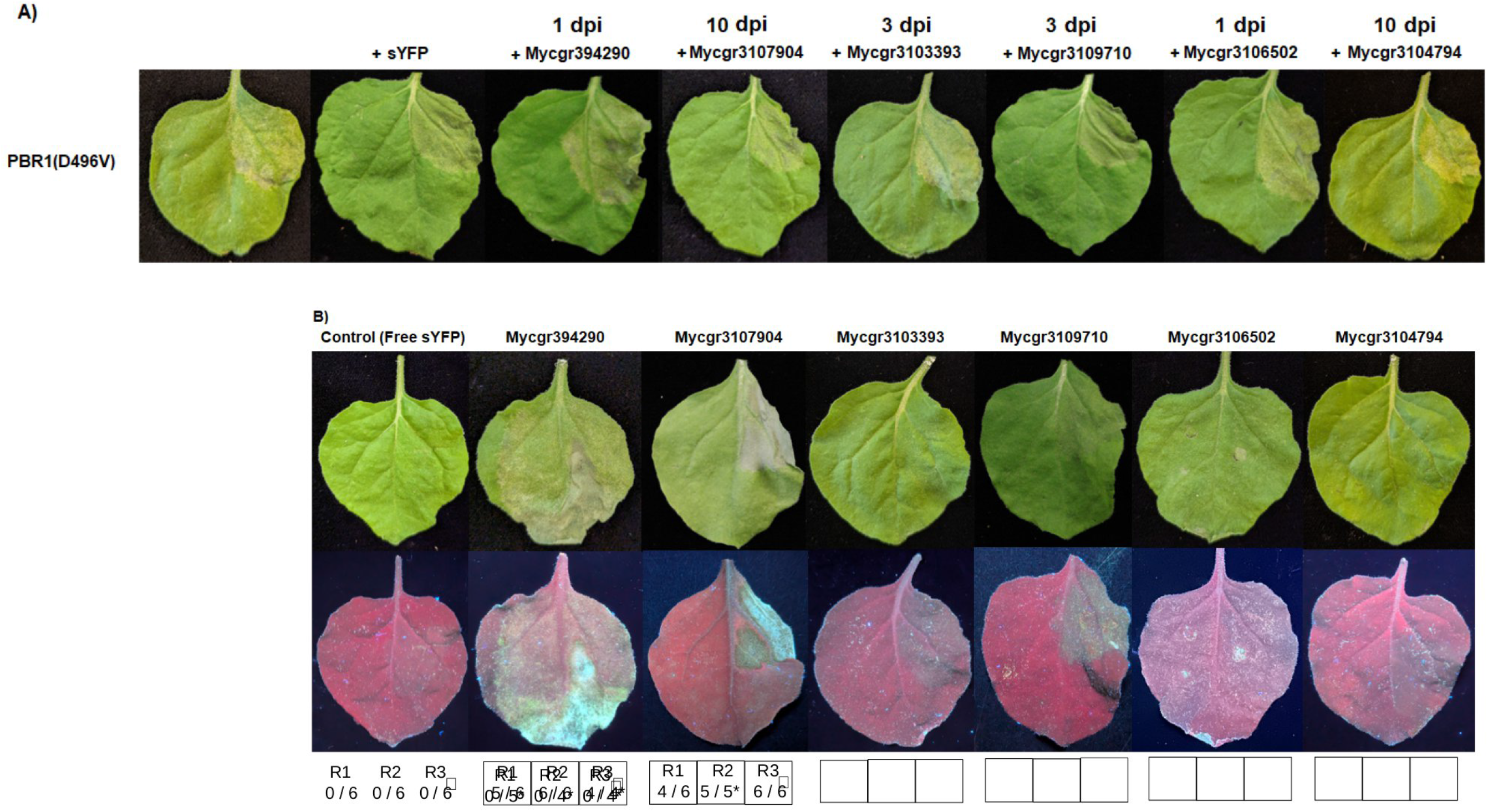
Responses of *Nicotiana benthamiana* leaves to infiltration with *Agrobacterium tumefaciens* express *tritici* effector proteins with their signal peptides. A) Suppression of PBR-1-induced cell death in *N. benthamian* infiltrated with *A. tumefaciens* expressing PBR-1(D496V) and either free sYFP as a control or a candidate effect peptide. B) Induction of necrosis caused by candidate effectors from *Z. tritici.* Leaves of *N. benthamiana* were in *tumefacien*s expressing an effector candidate with its signal peptide. Necrosis was observed at six days post infil circles indicate regions where cell death was observed. The squares at the bottom of the figure indicate the numb replicates (R1, R2, R3) and the quantification of response frequency (proportion of leaves that showed cell death Asterisks indicate replicates where fewer than six leaves in total were evaluated due to some of the leaves showi the infiltration. The top row is under normal light. The bottom row is under ultraviolet light. The day when each showed its highest level of expression in previous RNAseq experiments is indicated at the top of panel A with d post-inoculation.

### 2.5. Two candidate effectors induce cell death in *N. benthamiana*

*Agrobacterium*-mediated transient expression of candidate effectors Mycgr3107904 and Mycgr394290, including their signal peptides, triggered cell death in *N. benthamiana* leaves six days post infiltration (dpi) (Figure 4B). In contrast, no cell death was observed in plants infiltrated with the control treatment (free sYFP). Mycgr3107904 contains an Hce2 domain, which has been associated with cell death induction in effectors from other pathogens (Stergiopoulos et al., 2012; Zhang et al., 2019) and in the necrosis-inducing protein ZtNIP1 from *Z. tritici* (Ben M’Barek et al., 2015). These findings support a potential role for Mycgr3107904 in triggering cell death in *N. benthamiana*. For Mycgr394290, the observed induction of cell death is consistent with the trend toward increased ROS production when the effector is expressed with its signal peptide. However, this effect on ROS accumulation did not reach statistical significance, despite being observed across biological replicates. These observations suggest that Mycgr394290 may promote a ROS burst that contributes to the activation of cell death responses. Nevertheless, further experiments will be required to confirm this relationship and to establish the underlying mechanism.

## 3. DISCUSSION

These results provide new insights into the potential functional roles of candidate effectors expressed during *Z. tritici* infection of wheat. Our findings help clarify how specific effector proteins contribute to modulate host immunity and whether their activity aligns with distinct stages of disease progression.

Given the limited sequence similarity among fungal candidate effectors, we used AlphaFold v3.0 to predict the tertiary structures of 32 *Zymoseptoria tritici* effectors. This allowed us to identify confidently predicted regions and domains. Most amino- and carboxy-terminal regions showed low confidence scores (pLDDT < 50) (Figure 2A), consistent with intrinsic disorder rather than unpredicted structure (Ruff and Pappu, 2021). Intrinsically disordered regions evolve more rapidly than ordered regions and can adopt multiple conformations, which may facilitate effector translocation or interactions with host proteins (Marín, Uversky and Ott, 2013). Indeed, experimental and computational evidence increasingly supports the prevalence of disordered regions in plant pathogen effectors (Wulf et al., 2004; Rodgers et al., 2008; Seong and Krasileva, 2023; Chepsergon and Moleleki, 2023).

We performed structural comparisons using both the FoldSeek server (https://search.foldseek.com/search) (Van Kempen et al., 2024) and the DALI server (Holm, 2022) to assess whether highly expressed *Z. tritici* effectors share significant structural similarity or folding patterns with previously identified and characterized effectors. We found that candidate effectors Mycgr3111221 and Mycgr3105487 share structural similarity with the *F. fulva* LysM effector Ecp6. Their amino acid conservation predicts a functional similarity between these effectors and Ecp6, particularly in their likely abilities to bind chitin fragments and suppress chitin-triggered immune responses in host plants (de Jonge et al., 2010; Marshall et al., 2011; Sánchez-Vallet et al., 2020; Tian et al., 2021).

There are several possibilities for how effectors can function in host cells. Recognition of PAMPs or damage-associated molecular patterns (DAMPs) by PRRs triggers transient ROS production. This is one of the early signatures of cell surface immunity or basal immunity in plants (Jwa and Hwang, 2017; Torres, 2024). Alternatively, effectors can enhance ROS production by interacting directly or indirectly with intracellular NLR proteins, ultimately leading to HR-associated cell death (Greenberg and Yao, 2004; Jwa and Hwang, 2017). We hypothesize that high expression of the selected effector genes from *Z. tritici* during the early (1 and 3 DAI) and transition to necrotrophy (10 DAI) stages of infection may reflect roles in modulating ROS accumulation, potentially suppressing or altering immune signaling consistent with their expression timing. In fact, AlphaFold predictions revealed structural similarity that supports the functional hypotheses; for instance, Mycgr3107904 shares similarity with VmHEP1 from *V. mali,* an effector known to induce necrosis in *N. benthamiana* (Zhang et al., 2019), suggesting a role in enhancing ROS production or triggering cell death.

The *Z. tritici* effector Mycgr3109710, which shows elevated expression at 3 dpi, shares structural similarity with PRY1, a secreted glycoprotein from *Saccharomyces cerevisiae* that is known to bind and export cholesteryl acetate (Darwiche et al., 2016). PRY proteins belong to the cysteine-rich secretory, antigen 5, and pathogenesis-related 1 (CAP) protein superfamily (Han et al., 2023), whose members display diverse physiological functions, including roles in fungal virulence (Darwiche et al., 2016; Han et al., 2023). PRY1 binds sterols and associates with free fatty acids (FAs), promoting their export in *S. cerevisiae* (Darwiche et al., 2016). During plant immune responses, phospholipase activity remodels membrane lipids and releases FAs and signaling lipids such as phosphatidic acid (PA), which modulates ROS production through regulation of NADPH oxidases (Hou, Ufer and Bartels, 2016; Seth et al., 2024). Given the structural similarity between PRY1 and Mycgr3109710, and the reported function of PRY1 from *S. cerevisiae,* it is possible that Mycgr3109710 suppresses ROS accumulation by binding host lipids, potentially sequestering free FAs or PA and thereby dampening downstream immune signaling.

In several plant pathogens, PR-1–like (PR1-L) proteins have been shown to act as virulence determinants by directly manipulating host cellular processes. In *Ustilago maydis*, UmPR-1La binds plant-derived phenolics to induce hyphal-like growth and is proteolytically processed by the maize cysteine protease CatB3 at a conserved CNYD motif, releasing a CAP-derived peptide (CAPE)-like peptide that suppresses CAPE-primed immunity while avoiding cleavage of endogenous plant PR-1 proteins (Lin et al., 2023). Similarly, in *Cytospora chrysosperma*, the PR1-L proteins CcCAP1 and CcCAP2 interact with the host RNA polymerase II subunit (RPB11), with CcCAP1 proposed to stabilize this factor to modulate transcription and dampen PR1 gene expression (Han et al., 2026). Beyond host manipulation, deletion of CcCAP2 and CcCAP3 impaired filamentous growth, implicating PR1-L proteins in fundamental fungal processes (Han et al., 2026). The conservation of the CAP domain across PR-1 proteins points to a shared mechanistic basis for these functions (Choudhary and Schneiter, 2012; Han et al., 2023). Structural analyses of CAP family members have revealed a characteristic α–β–α sandwich fold stabilized by disulfide bonds linking the central β sheet to C-terminal helices (Fernández et al., 1997; Han et al., 2023). Consistent with this architecture, the predicted tertiary structure of Mycgr3109710 adopts a similar α–β–α fold configuration, with disulfide bonds likely contributing to its structural stability.

Foldseek and DALI structural similiarity analyses also revealed that the candidate effector Mycgr3107904 shares structural similarity with Hce2 domain-containing effectors from other pathogenic fungi, including VmHEP1 from *Valsa mali* (Figure 2B). Among five Hce2 effectors encoded in the *V. mali* genome, VmHEP1 alone triggered cell death in *N. benthamiana* and contributed to virulence on apple. A double knockout of VmHEP1 and the related effector VmHEP2 significantly reduced pathogenicity, despite only 36% sequence identity between them (Zhang et al., 2019). Sequence divergence and complementary expression patterns between VmHEP1 and VmHEP2 revealed by qRT-PCR suggested that *V. mali* may evade host recognition and maintain virulence by compensating for the suppression of one effector through upregulation of the other (Zhang et al., 2019). This strategy also may be relevant for *Z. tritici*, which encodes three Hce2 domain-containing effectors: Mycgr3107904, Mycgr3111636, and Mycgr3104404 (Stergiopoulos et al., 2010, 2012). These proteins, ranging from 159–179 amino acids and predicted to be secreted, share 43.7–58.0% sequence identity, with Mycgr3107904 and Mycgr3111636 showing the highest similarity (58.0%) and strong expression at 10 DAI (Supplementary table 1).

Due to the sequence identity of Mycgr3107904 and the other two Hce domain-containing effectors, we were curious to see if they also share structural similarity. Using the RSCB pairwise comparison tool, we found that the structural alignment between Mycgr3107904 and Mycgr3111636 generated a TM score of 0.77 and a RMSD of 2.21, revealing structural similarity (Supplementary figure 2). Meanwhile, structural alignments between Mycgr3104404 and Mycgr3111636 and Mycgr3107904 generated TM scores of 0.57 and 0. 63 and RMSD scores of 3.31 and 3.72, respectively (Supplementary figure 2). This suggests that *Z. tritici* might employ sequence variation and co-expression of these effectors to avoid host detection, similar to *V. mali*. Moreover, in *Z. tritici*, a fourth Hce2 domain-containing effector, named ZtNIP1, was identified through crude culture filtrates and liquid chromatography-mass spectrometry (Ben M’Barek et al., 2015). ZtNIP1 induces necrosis in wheat cultivars, aligning with our observation that Mycgr3107904 triggers cell death when transiently expressed in *N. benthamiana* (Figure 4B). Taken together, these observations suggest that the *Z. tritici* candidate effector Mycgr3107904 is capable of triggering cell death when expressed in *N. benthamiana,* likely reflecting immune recognition in this non-host system rather than its native activity in wheat. While this response may indicate that Mycgr3107904 can interact with plant immune components and potentially influence defense-associated programmed cell death pathways, its contribution to virulence during *Z. tritici* infection of wheat remains to be determined. It also is plausible that Mycgr3107904 acts either independently or in coordination with other Hce2 domain-containing effectors to directly promote cell death or modulate host defenses. However, further functional analyses in the host context are required to test this hypothesis.

While our study utilized transient expression of epitope-tagged effector proteins in a model plant species, the consistent attenuation of chitin-induced ROS production by some candidate effectors expressed with their signal peptides compared with the expression without their signal peptides supports their classification as secreted, functional effectors, and provides evidence that the presence of the signal peptide is required for the suppression of cell-surface immunity responses.

The observed pattern of certain effectors to either suppress or enhance ROS production when transiently expressed inside the cell (with no signal peptide) may reflect non-specific effects or technical artifacts associated with heterologous overexpression and mislocalization, rather than genuine biological function. However, it is worth considering the possibility of a potential intracellular function. Although effector translocation from the apoplast into host cells has not yet been demonstrated in *Z. tritici* (Meile et al., 2024), this possibility should be further explored. Notably, extracellular vesicles (EVs) have been isolated from *Z. tritici* culture filtrates and characterized as membrane-bound particles containing a diverse protein cargo similar to those reported in EVs from other pathogenic fungi (Hill and Solomon, 2020). While EVs in human pathogens have been implicated in virulence and host modulation, their contribution to plant-fungal interactions and effector delivery in phytopathogenic fungi, including *Z. tritici*, remains unclear (Wang et al., 2023). Therefore, while an intracellular role cannot be excluded, these results should be interpreted with caution and will require validation. Future analyses examining whether secreted effectors can access host intracellular compartments, potentially via EV-associated or other noncanonical secretion pathways, will be important for understanding effector function during infection.

The functional assay results suggest that Mycgr3109710 plays a localization-dependent role in host-pathogen interactions, as it significantly suppresses ROS production when the signal peptide is present. Its structural similarity to PRY1 and other CAP domain-containing fungal effectors, annotated as PR-1-like, indicates a possible role in pathogenicity. One hypothesis is that Mycgr3109710 binds plant sterols, disrupting membrane integrity, similar to the antifungal action of plant PR-1 proteins (Gamir et al., 2017; Kazan and Gardiner, 2017). In fact, the localization of PR-1 proteins from plants, whether extracellular or vacuolar, has been shown to influence their role in plant basal defense (Kazan and Gardiner, 2017). This suggests that *Z. tritici* may exploit the localization of Mycgr3109710 to fine tune host immune responses and attenuate ROS to its advantage. This strategy may extend to other PR-1-like effectors such as Mycgr3103393, which only suppresses ROS when expressed with its signal peptide.

The likely role of candidate effector Mycgr394290 was hard to predict from the experimental results. The observed pattern with its signal peptide showed that it increases ROS production compared to free sYFP, although this effect was not statistically significant. However, expressing Mycgr394290 with its signal peptide induces cell death in *N. benthamiana* (Figure 4B). Effectors can interact directly or indirectly with NLR proteins to amplify ROS production and trigger HR cell death, highlighting the crucial role of ROS in NLR-mediated immunity (Greenberg and Yao, 2004; Jwa and Hwang, 2017). For instance, the *F. fulva* effector Avr2, upon binding to the extracellular cysteine protease Rcr3 in tomato, induces both ROS accumulation and cell death (Shabab et al., 2008). Mycgr394290 appears to be a species-specific effector in *Z. tritici*, as it lacks any known domains based on an InterPro search. Its tertiary structure prediction includes regions with low to very low confidence, and no structural homologs were identified using FoldSeek. Additionally, Mycgr394290 is present in the same region of chromosome 7 in all 18 *Z. tritici* isolates, even though it was not predicted as a protein in 15 of these isolates. This candidate effector also appears to be disrupted in the three European isolates 3D7, 1A5, and 1E4, where it likely does not encode a full-length functional protein. The annotated short protein (∼60 aa) to which it aligns in those three European isolates is therefore most likely a truncated remnant of the original gene rather than a bona fide functional effector. This aligns with previous findings in the reference genome of *Z. tritici*, where chromosome 7 exhibits high gene density but predominantly encodes species-specific genes (Feurtey et al., 2020), such as candidate effector gene Mycgr394290. Additionally, chromosome 7 shares interesting characteristics with accessory chromosomes, such as reduced gene expression and lower H3K4me2 methylation levels. Studies across *Z. tritici* isolates have shown that most predicted effector genes are located on core chromosomes, with only a few found on accessory chromosomes (Feurtey et al., 2020).

Overall, our observations suggest that the presence of a signal peptide significantly influences the functionality of candidate effectors in *Z. tritici*. Supporting our findings, Thynne et al. (2024) demonstrated that Mycgr3107904 (referred to as Zt_2_242 in their study) attenuates flg22-induced ROS production when transiently expressed with its signal peptide in *N. benthamiana* leaves. Similarly, for Mycgr3111221, an extensively characterized effector in *Z. tritici* called Mg3LysM by Marshall et al. (2011), the results of our ROS assays align with those of previous studies showing that this effector suppresses chitin-induced ROS production by binding to chitin fragments (Marshall et al., 2011; Lee et al., 2014). This interaction not only inhibits ROS production but also was confirmed to protect fungal hyphae from degradation by plant-derived hydrolytic enzymes (Marshall et al., 2011).

A limitation of the current study is that effector secretion was assessed using total leaf protein extracts (Figure 3B), rather than isolated apoplastic fluids. While this approach confirms effector accumulation upon expression, it does not provide direct subcellular resolution of secretion to the apoplastic space. Apoplastic fluid isolation through vacuum infiltration-centrifugation, followed by immunoblot detection and verification of fraction purity using cytoplasmic contamination markers (Joosten, 2012; Hu et al., 2021), would offer more definitive evidence that signal peptide-bearing effectors are actively secreted and functional at the host-pathogen interface. This represents a priority experiment for future work, where apoplastic proteomics combined with effector-specific detection could be employed to validate the localization-dependent activities described here and to further dissect the spatiotemporal dynamics of effector deployment during infection.

In our study, we also used PBR1(D496V)-mediated cell death as a proxy to explore whether *Z. tritici* candidate effectors are capable of suppressing NLR-associated immune signaling. PBR1 is a well-characterized NLR protein from barley that recognizes the protease activity of the bacterial effector AvrPphB from *P. syringae* (Carter et al., 2019). The *PBR1* (D496V) allele carries a mutation in the conserved MHD (Methionine-Histidine-Aspartate) motif (MHD → MHV), resulting in constitutive activation of the protein. When PBR1 (D496V) : sYFP is expressed transiently in *N. benthamiana*, it triggers macroscopic cell death within 24 hours following dexamethasone induction (Jaiswal et al., 2023). Additionally, Carter et al. (2019) demonstrated that wheat cultivars also recognize AvrPphB protease activity and possess two putative orthologs of *Pbr1*, suggesting a conserved role for PBR1 in immune responses across cereal crops. Therefore, the PBR1 (D496V) system provides a sensitive platform to detect suppression of downstream NLR signaling.

Testing of PBR1 is important because pathogen effectors can suppress NLR-triggered immunity indirectly, for example by targeting NLR-associated signaling components, helper NLRs, oligomerization processes, or downstream signaling events, rather than by interacting directly with the receptor itself (Wu and Derevnina, 2023). Consistent with this concept, some effectors from non-bacterial pathogens have been shown to suppress intracellular NLR signaling despite being apoplastic, including the nematode effectors *Gr*UBCEP12 and *Gr*EXPB2 which inhibited cell death induced by the autoactive mutant of the NLR Rx (AtRx) in *N. benthamiana* and *N. tabacum* leaves, proving that apoplastic effector proteins can affect intra-cellular signaling pathways (Ali et al., 2015). Other effectors also have been shown to suppress NLR-mediated responses through intermediary host proteins, including the oomycete effector AVRcap1b, which suppresses NLR-dependent cell death via the membrane trafficking protein NbTOL9a (Derevnina et al., 2021a). Together, these observations provide a rationale for using PBR1(D496V) as a generalized assay to probe the potential of *Z. tritici* effectors to suppress NLR-dependent cell death signaling.

Contrary to our expectations, none of the seven tested effectors suppressed the cell death triggered by this NLR (Figure 4A). PBR1 was selected as a well-characterized receptor that provides a robust and reproducible cell death readout in *N. benthamiana*, enabling us to assess effector-mediated interference with immune signaling at a general level. This assay does not imply a direct biological interaction between PBR1 and *Z. tritici* effectors in wheat; rather, it tests whether these effectors can block NLR-dependent cell death once signaling has been activated.

Although the tested effectors are predicted to be secreted or apoplastic, accumulating evidence suggests that apoplastic effectors can be internalized into host cells or exert indirect effects on intracellular immune signaling via metabolic or regulatory perturbations (Ali et al., 2015; Derevnina et al., 2021). Nonetheless, the use of PBR1, an NLR known to guard PBS1 against bacterial proteases, has limitations when extrapolating to the wheat - *Z. tritici* pathosystem, particularly given the lack of currently characterized NLRs conferring resistance to *Z. tritici* (Saintenac et al., 2021; Hafeez et al., 2025). Future studies using wild-type PBR1 activated by its cognate effector (AvrPphB), or preferably wheat or barley NLR candidates such as the mildew resistance locus A (MLA) (Lu et al., 2016), as well as *Z. tritici* effectors known to induce necrosis in *N. benthamiana* such as effector Zt6 (Kettles et al., 2018), will provide a more biologically relevant framework to test effector-mediated suppression of host immunity.

Our analyses relied on a non-host plant model system, where effectors were expressed with or without their putative signal peptides, which may have influenced their proper delivery into the apoplastic space. To confirm their correct processing, future analyses should express *Z. tritici* candidate effectors in *N. benthamiana* using native *N. benthamiana* signal peptides. Notably, *N. benthamiana* is an effective model for analyzing effector functionality, allowing efficient DNA integration and expression after infiltration with *Agrobacterium*. At least two studies have successfully used this heterologous system to characterize candidate effectors of *Z. tritici* (Kettles et al., 2017; Thynne et al., 2024). However, an important consideration is that the occasionally inconsistent ROS phenotypes observed in this study may stem from the use of *N. benthamiana* as a heterologous system. This non-host plant likely harbors immune receptors or signaling components that differ from those present in wheat, which may result in apparent ROS suppression or activation that does not reflect the native activity of these effectors in their natural host.

Future research should also determine whether the immune manipulation activities of Mycgr3109710, Mycgr3107904, Mycgr394290, and other candidate effectors are reproducible in wheat. This could be achieved using alternative delivery systems, such as *Pseudomonas fluorescens* strain EtHAn, which was successfully employed to introduce the *Puccinia striiformis* effector Shr7 into wheat cells, revealing its immune-suppressive function (Ramachandran et al., 2017). Exploring similar bacterial systems could help assess whether the candidate effectors studied here exert the same immune-modulating effects in their natural host.

## 4. EXPERIMENTAL PROCEDURES

### 4.1. Selection of transcriptionally induced candidate effectors

Gene expression data for candidate effectors at the asymptomatic stages of infection (1-, 3-, and 6- days after inoculation (DAI), and at the transition to the necrotrophic stage (10 DAI) was obtained from Gomez-Gutierrez et al. (2025). From this dataset, we first selected 32 candidate effector genes that were differentially expressed in the susceptible interaction compared to a non-host interaction (Gomez-Gutierrez et al., 2025) (Supplementary Table 1). Signal peptides were confirmed using SignalP v6.0 (Teufel et al., 2022) and Phobius (v1.01) (Käll, Krogh and Sonnhammer, 2004) by applying a probability threshold of 0.9. Transmembrane domains were assessed and predicted to be absent using Phobius (v1.01) (Käll, Krogh and Sonnhammer, 2004) and DeepTMHMM (v1.0.24) (Hallgren et al., 2022). EffectorP (v3.0) (Sperschneider and Dodds, 2022) was employed to identify *Z. tritici* proteins with effector-like sequence features.

Subsequently, we narrowed the focus to seven candidate effectors that have a higher expression as demonstrated by their normalized counts in the replicates of the susceptible interaction in either the early stages of infection or the transition to the necrotrophic stage for functional characterization (Table 1).

**TABLE 1.**
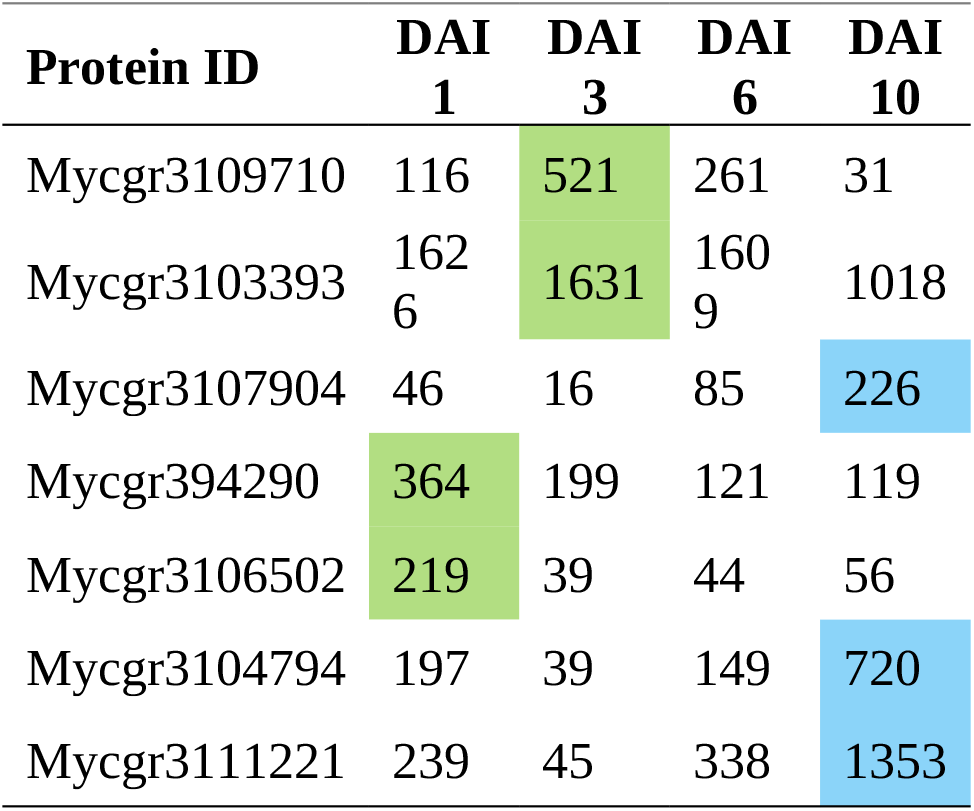
Normalized count values for a subset of candidate effectors in the genome of *Zymoseptoria tritici* that are highly expressed in either the early stage of infection (1 – 3 days after inoculation (DAI)) or at the transition to the necrotrophic stage (10 DAI), with the highest counts for each effector highlighted in green color for 1 and 3 DAI, and in blue color for 10 DAI.

### 4.2. Conservation of candidate effectors in *Zymoseptoria* isolates

We assessed the conservation of 32 candidate effectors (Supplementary Table 1) across *Z. tritici* isolates and the sister species *Z. pseudotritici* (Stukenbrock et al., 2012)*, Z. brevis* (Grandaubert, Bhattacharyya and Stukenbrock, 2015), and *Z. ardabiliae* (Stukenbrock et al., 2012). Predicted protein sequences and genome assemblies for *Z. tritici* IPO323 and its sister species were retrieved from the JGI MycoCosm portal (https://mycocosm.jgi.doe.gov/mycocosm/), while those for 18 additional *Z. tritici* isolates were obtained from the GitHub repository of the Laboratory of Evolutionary Genetics of Dr. Daniel Croll at the University of Neuchâtel (https://github.com/crolllab) (Badet et al., 2020). Using the 32 candidate effector sequences from the reference strain IPO323 as queries, we performed BLASTp searches (E value ≤ 10^⁻10^) against a proteome database of 21 *Zymoseptoria* isolates. The nucleotide sequences of the 32 candidate effectors were retrieved from Ensembl Fungi Biomart (Kinsella et al., 2011) and tBLASTx searches (E value ≤ 10⁻Z⁰) were conducted against a nucleotide database containing the genome scaffolds of all 22 isolates including IPO323. The databases for BLAST were generated using the “makeblastdb" function from BLAST (Boratyn et al., 2013).

### 4.3. Prediction of protein structures using AlphaFold3

The structures of 32 candidate effector sequences were predicted using AlphaFold v3.0 (Abramson et al., 2024). Signal peptides were removed prior to modeling by processing the sequences with SignalP v6.0 (Teufel et al., 2022). AlphaFold v3.0 generated five models for each candidate effector, ranked from 0 to 4 based on their confidence scores, and the corresponding PAE plots were downloaded as well The model with the highest Predicted Local Distance Difference Test (pLDDT) score (model_0.cif) was selected for further analysis. A pLDDT score above 90 indicates high accuracy in both backbone and side chain predictions, while scores between 70 and 90 suggest accurate backbone predictions but potential inaccuracies in side chain placement (Jumper et al., 2021). This selected CIF file was used as the query input for the Foldseek web server (https://search.foldseek.com/search) (Van Kempen et al., 2024) and the DALI server (Holm, 2022) to identify proteins with potential structural similarity to the candidate effectors. The search databases in Foldseek were under default settings including AFDB/Proteome-v4, AFDB50, AFDB/Swiss-Prot-v4, CATH50-v4.3.0, GMGCL-2204, MGnify-ESM30-v1, and PDB100. The DALI server compares the input query structure against the Protein Data Bank and generated a list of hits ranked by Z-score. Foldseek generated a list of hits ranked by structural similarity scores, which included the alignment score (TM-score) and the Root Mean Square Deviation (RMSD) of aligned regions. Results were downloaded for further visualization and analysis. Predicted structures and structural alignments between the query protein and identified homologs were visualized using ChimeraX v1.7 (Meng et al., 2023).

AlphaFold v3.0 was also used to predict the tertiary structure of VmHEP1 effector protein from *Valsa mali* (Zhang et al., 2019), a likely structural homolog of candidate effector Mycgr3107904 in *Z. tritici.* We performed pairwise structural alignment of the tertiary structure of VmHEP1 against the predicted structure of Mycgr3107904 using the RCSB PDB tool (https://www.rcsb.org/alignment) (Bittrich et al., 2024).

### 4.4. Plant growth conditions

*Nicotiana benthamiana* was cultivated in Berger Seed Germination and Propagation Mix supplemented with Osmocote slow-release fertilizer (14-14-14). Plants were kept in a controlled environment with a 16-hour light and 8-hour dark cycle, with temperatures set to 24°C during the light phase and 22°C during the dark phase, and maintained at 60% humidity. The average light intensity measured at plant height was 130 µmol/mZ/s.

### 4.5. Generation of plant expression constructs

Plant expression constructs were generated for four candidate effectors that exhibited significant expression at early infection stages (1 and 3 DAI) and for three candidate effectors that have higher expression during the transition to the necrotrophic phase (10 DAI) using a modified multisite Gateway cloning strategy (Qi, DeYoung and Innes, 2012; Helm et al., 2022).

Constructs for these effectors were generated with and without their native signal peptides to conduct functional characterization experiments for both versions of the effectors. Open reading frames (ORFs) of *Zymoseptoria tritici* candidate effectors, both including and excluding their predicted signal peptides, were synthesized by Azenta Life Sciences and inserted into the pUC57 plasmid as Gateway-compatible donor clones (pDONR[L1-L4]:ZtEC).

To generate ZtEC:sYFP fusions, pDONR(L1-L4):ZtEC constructs were combined with pBSDONR(L4r-L2):sYFP (Qi, DeYoung and Innes, 2012; Helm et al., 2022) and recombined into the plant expression plasmid pEG100 (Earley et al., 2006) using LR Clonase II (Invitrogen), following the manufacturer’s protocol. All constructs were verified by sequencing before use in transient expression assays.

### 4.6. Transient protein expression in *N. benthamiana*

Agrobacterium-mediated transient expression was performed as described previously (Helm et al., 2022) with modifications. *Agrobacterium tumefaciens* strain GV3101 was transformed with the constructs and cultured in Luria-Bertani (LB) broth supplemented with gentamicin and kanamycin at 30°C for two days. Transformed colonies were inoculated into 5 mL of liquid LB with antibiotics and were shaken overnight at 30°C. The following day, bacterial cells were pelleted, resuspended in 10 mM MgCl₂, and adjusted to an OD₆₀₀ of 0.6. The suspensions were incubated with 100 µM acetosyringone for 3–4 hours at room temperature before infiltration into the abaxial surface of 3–4-week-old *Nicotiana benthamiana* leaves using a needleless syringe.

### 4.7. Measurement of Reactive Oxygen Species (ROS) in *N. benthamiana*

We followed the luminol-based protocol for the ROS assay as described by Jaiswal et al. (2022). Three-week-old *N. benthamiana* leaves were used for the experiment, infiltrated with *A. tumefaciens* carrying constructs that expressed either free sYFP or candidate effector-fluorescent protein fusions. Two days post infiltration (dpi), 12 leaf discs (5 mm diameter) per treatment were harvested using a cork borer and placed in a petri dish with distilled water. The discs were washed three times with sterile water, and then the petri dishes were placed in the dark under aluminum foil overnight. The following day, the distilled water was replaced, and the 12 leaf discs were placed in a white-bottomed, 96-well OptiPlateTM microplate (Perkin Elmer) (free sYFP leaf discs were placed in wells in row A, and discs of two different candidate effectors were placed in rows B and C), in 200 μL of Milli-Q water. Immediately before reading, Mili-Q water was removed and leaf discs in rows A, B and C were treated with chitin elicitation solution (luminol [30 μg/mL], horseradish peroxidase [20 μg/mL], chitin [hexamer, 5 μg/mL], and nuclease-free water). The relative luminescence accumulation (RLU) was measured over 40 minutes in a microplate reader (Tecan Infinite M200 Pro). These experiments were performed in three to six independent biological replicates. In some assays, neither the sYFP control nor effector-expressing leaf discs exhibited the expected kinetic profile of chitin-induced ROS production; in such cases, additional independent biological replicates were conducted to ensure the kinetic profile of chitin-induced ROS production was observed and only those biological replicates in which both the sYFP control and the effector-expressing leaf discs exhibited the expected kinetic profile were included in the statistical analysis. The average total RLU from all of the leaf discs (technical replicates) for each effector in each ROS assay was calculated and normalized by dividing the average total luminescence of effector-expressing leaf discs to the average of the leaf discs expressing the negative control free sYFP fluorescent protein. Data from three to four biological replicates for each effector was included to conduct a one-sample t-test (p-value = 0.05) and was used to represent the results (Figure 3A).

### 4.8. Necrosis induction assay

To identify putative effectors that induce cell death, *A. tumefaciens* cultures carrying constructs of candidate effectors with their native signal peptides were adjusted to an OD_600_ of 0.6 and infiltrated into defined areas of individual leaves of 3- to 4-week-old *Nicotiana benthamiana* plants. Each effector was infiltrated into at least two leaves per plant, with a total of three plants from the same pot. Cell death responses were assessed six dpi. As a control, *A. tumefaciens* carrying a construct expressing only the free sYFP tag was infiltrated in parallel. The experiment was performed three times.

### 4.9. Cell-death suppression assay

To identify putative effectors capable of suppressing cell death, we used a dexamethasone (DEX)-inducible construct expressing PBR1 (D496V) : sYFP (Jaiswal et al., 2023). PBR1, first described by Carter et al. (2019), is an NLR protein from barley that is cleaved by cysteine protease AvrPphB from *Pseudomonas syringae* (Carter et al., 2019). The *PBR1* (D496V) allele carries a mutation in the conserved Methionine-Histidine-Aspartate (MHD) motif, where the aspartate (D) residue at position 496 is substituted with valine (V), altering the motif from MHD to MHV and resulting in constitutive activation. Transient expression of PBR1 (D496V) : sYFP in *N. benthamiana* induces macroscopic cell death within 24 hours of dexamethasone treatment (Jaiswal et al., 2023).

To assess whether candidate effectors from *Z. tritici* suppress this response, we co-infiltrated *A. tumefaciens* suspensions carrying PBR1 (D496V) : sYFP (OD_600_ = 0.1) with those carrying individual effector constructs including their native signal peptides (OD_600_ = 0.6). The suspensions were mixed in equal volumes with 100 µM acetosyringone and incubated at room temperature for 3–4 hours before co-infiltration into the abaxial surface of 3- to 4-week-old *N. benthamiana* leaves. As control treatments, *A. tumefaciens* carrying a construct expressing PBR1(D496V) : sYFP and *A. tumefaciens* carrying a construct expressing free sYFP were co-infiltrated. Infiltrated leaves were sprayed with 400 mL of 50 µM DEX solution prepared from a 25 mM DEX stock two days after co-infiltration. Cell death was evaluated one day after spraying the leaves with DEX.

### 4.10. Immunoblot analyses

Immunoblotting was performed as described previously (Helm et al., 2022; Jaiswal et al., 2023) with modifications. *N. benthamiana* leaf discs (0.15 g) were collected 48 hours post infiltration, flash frozen, and homogenized in protein extraction buffer. Extracts were centrifuged, and the supernatant was mixed with 4, Laemmli buffer and beta-mercaptoethanol. Proteins were separated on a 4–20% Tris-glycine stain-free gel, transferred to a nitrocellulose membrane, and blocked in 5% skim milk. Recombinant sYFP-tagged proteins were detected using horseradish peroxidase-conjugated anti-GFP antibody (1:5,000) and visualized with Supersignal West Femto substrate using an ImageQuant 500 system.

## ACKNOWLEDGMENTS

The authors thank Drs. Terri Cameron and Ian Thompson (USDA-ARS, Crop Production and Pest Control Research Unit), Dr. Raksha Singh, and Jessica Cavaletto for technical assistance, scientific advice and insightful discussions. The funding bodies had no role in designing the experiments, collecting the data, or writing the manuscript. All opinions expressed in this paper are the authors’ and do not necessarily reflect the policies and views of USDA. USDA is an equal opportunity provider and employer.

## DATA AVAILABILITY STATEMENT

The data that support the findings of this study are openly available in Zenodo at https://doi.org/10.5281/zenodo.15477557.

## FUNDING

This research was funded by the United States Department of Agriculture, Agricultural Research Service (USDA-ARS) research project 5020-21220-014-00D.TABLES

**Supplementary Table 1** Normalized count values for thirty-two candidate effector genes from *Zymoseptoria tritici* that are highly expressed in either the asymptomatic stage of infection or during the transition to the necrotrophic stage, with the highest count in bold. DAI refers to days after inoculation of *Z. tritici*.

**Supplementary Table 2** Structural prediction and homology search results for candidate effectors of *Zymoseptoria tritici*. AlphaFold3-predicted structures were generated for 32 candidate effectors and evaluated using the predicted Template Modeling score (pTM) as a measure of overall model confidence, where values closer to 1.0 indicate higher structural reliability. Structural similarity searches were performed using FoldSeek and the Distance-matrix ALIgnment (DALI) server to identify proteins with similar three-dimensional folds in publicly available structure databases. For each candidate effector, the table reports the best FoldSeek match with its corresponding E value (where lower values indicate stronger structural similarity), and the best DALI match with its corresponding Z score (where values above 2 are generally considered significant). Candidate effectors with no significant structural homologs are indicated as "No match".

**Supplementary Figure 1.** Structural analysis of predicted effector proteins. **A.** PAE plots generated by Alphafold. **B.** Structural alignment of effector proteins using FoldMason. Effectors were grouped based on multiple structural alignment performed with FoldMason MSA (https://search.foldseek.com/foldmason).

**Supplementary Figure 2** Structural alignments of the Hce domain-containing effectors Mycgr3107904, Mycgr3111636 and Mycgr3104404.

