## Supplementary Figure 1 for "Assessing *Zymoseptoria tritici* Candidate Effectors in the Non-Host *Nicotiana benthamiana* Reveals Immune Recognition and Potential Host-Independent Activities"

**A)**

Mycgr382936

Mycgr379161

Mycgr346866

Mycgr339947

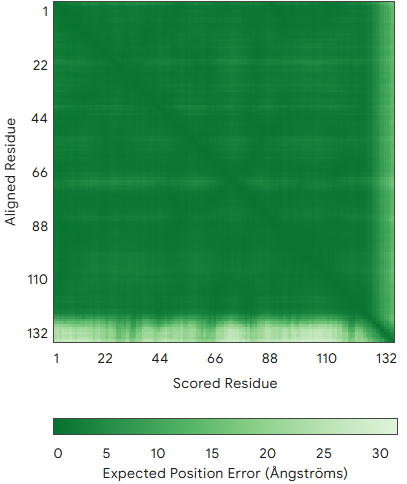

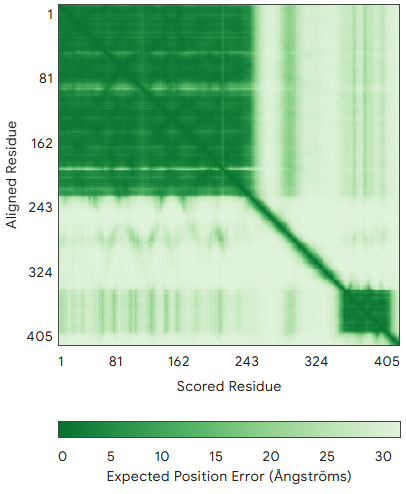

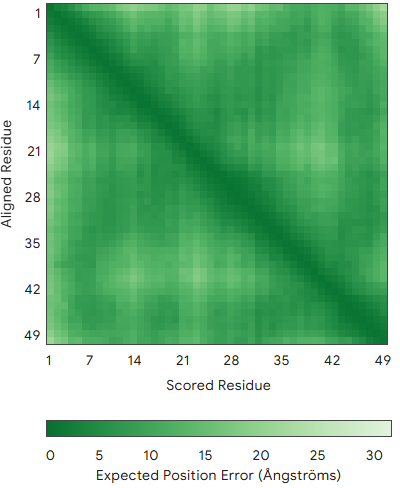

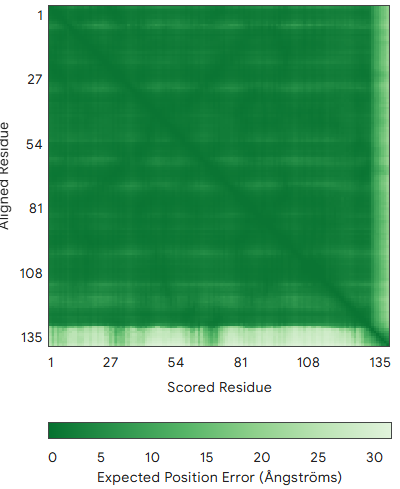

Mycgr3101652

Mycgr399161

Mycgr394290

Mycgr383064

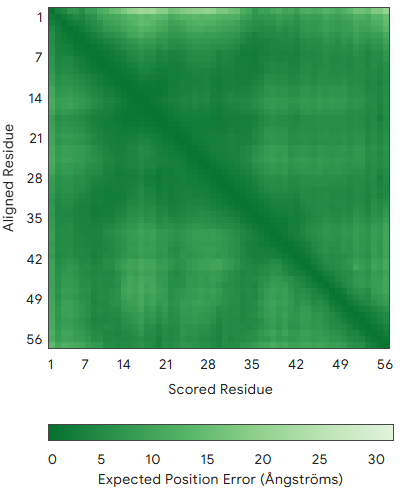

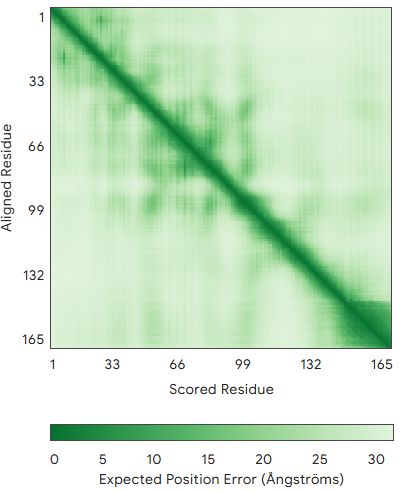

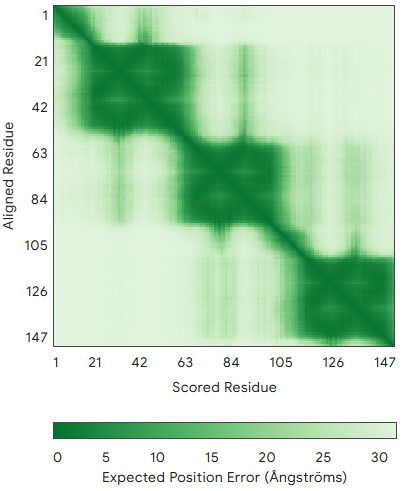

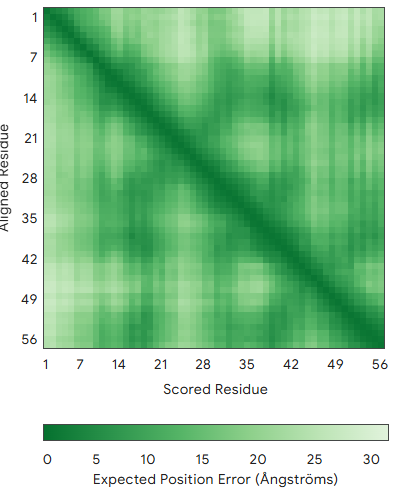

Mycgr3102792

Mycgr3102849

Mycgr3102996

Mycgr3103393

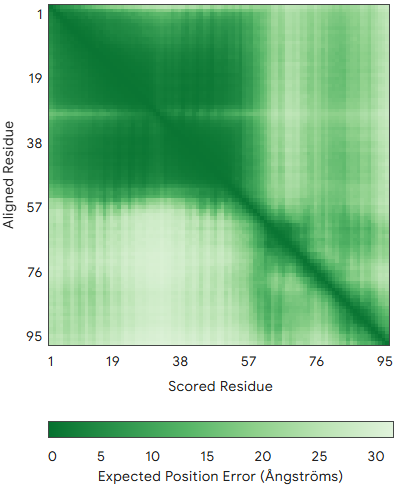

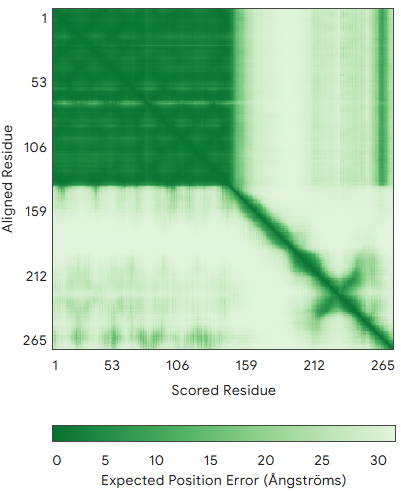

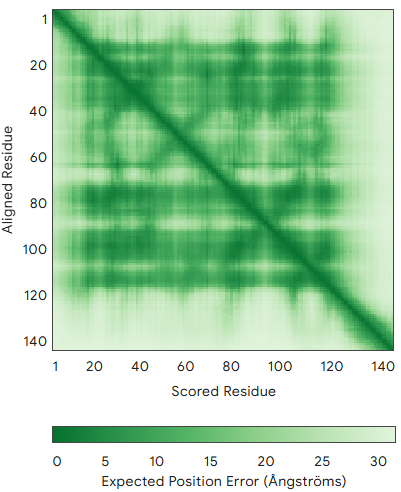

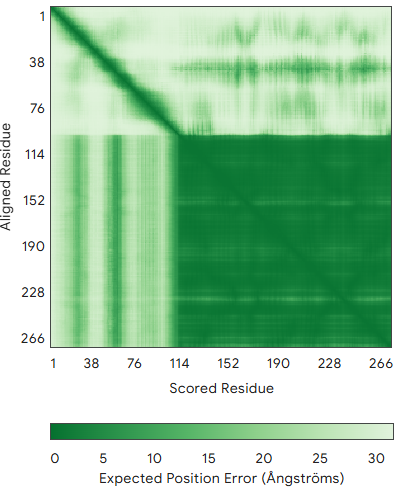

Mycgr3105182

Mycgr3104794

Mycgr3104697

Mycgr3104009

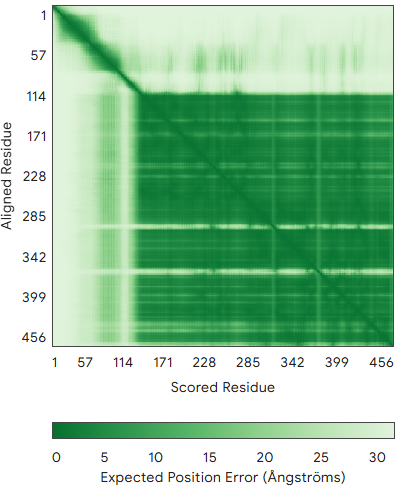

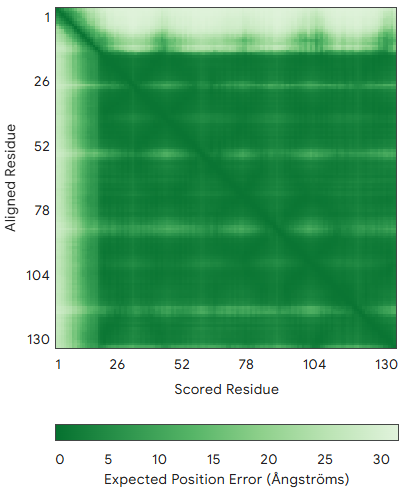

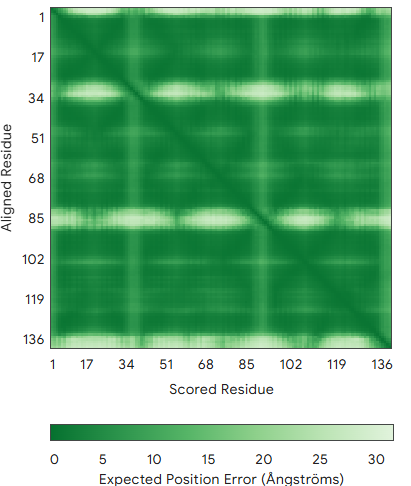

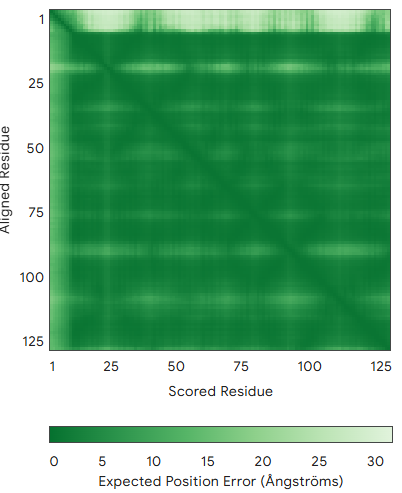

Mycgr3105223

Mycgr3105487

Mycgr3106106

Mycgr3106125

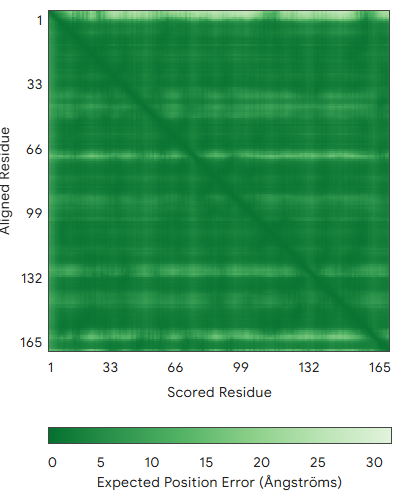

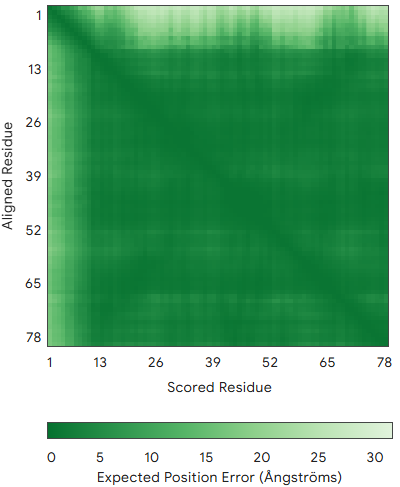

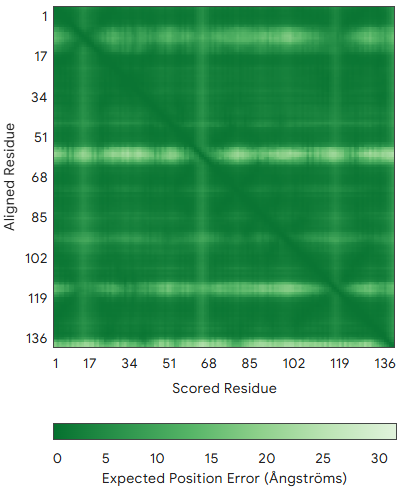

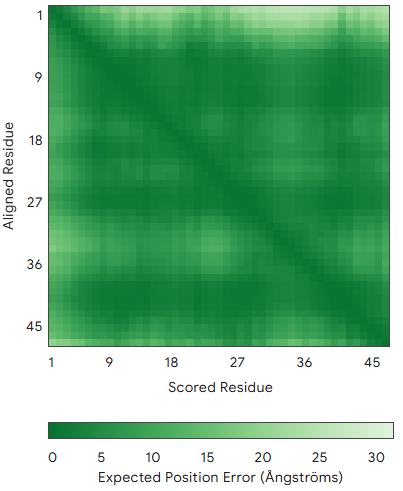

Mycgr3106456

Mycgr3107904

Mycgr3106502

Mycgr3106329

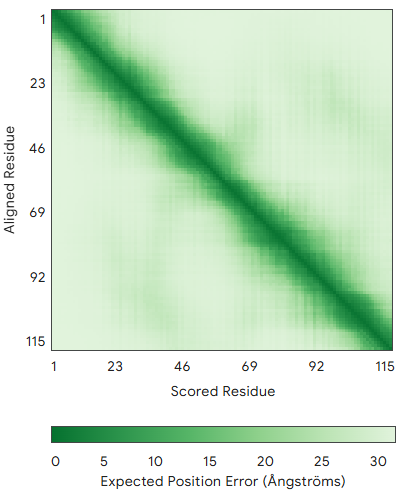

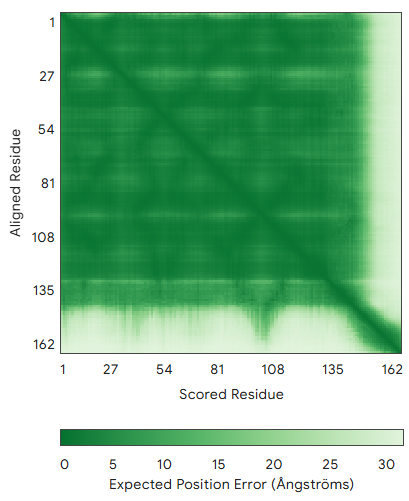

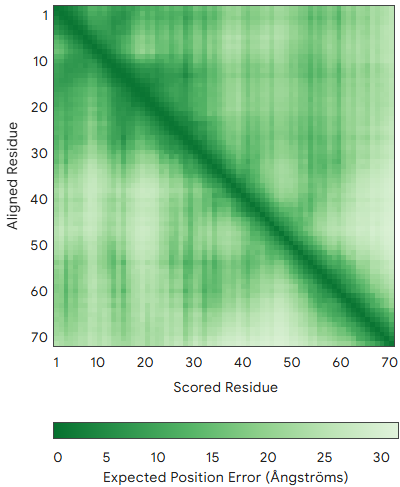

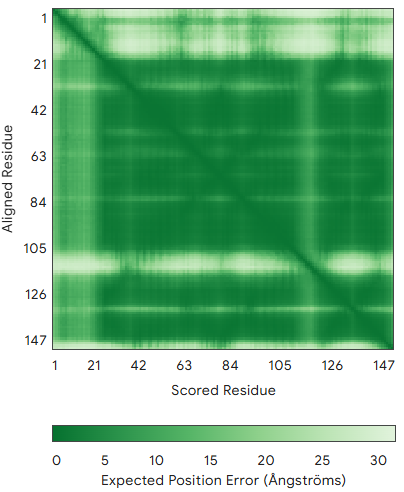

Mycgr3109137

Mycgr3109710

Mycgr3109991

Mycgr3110220

Mycgr3110887

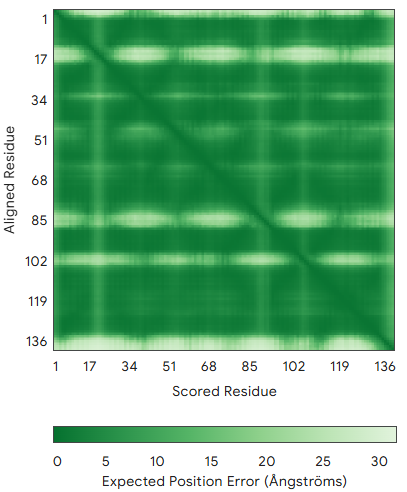

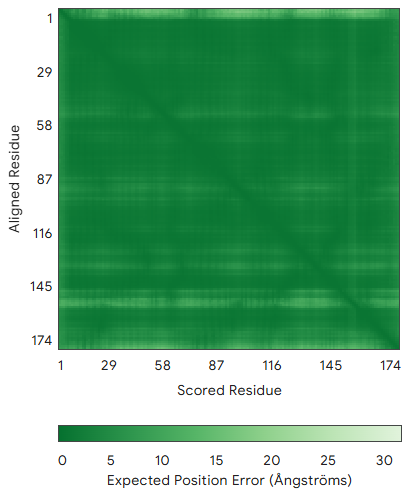

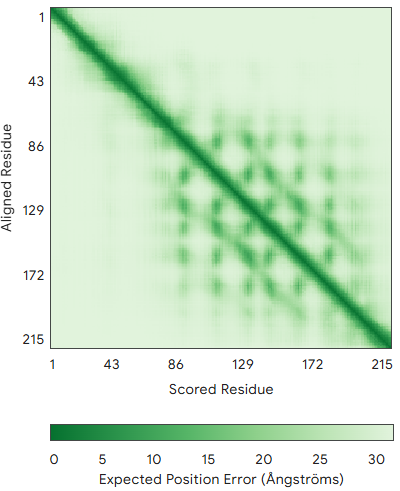

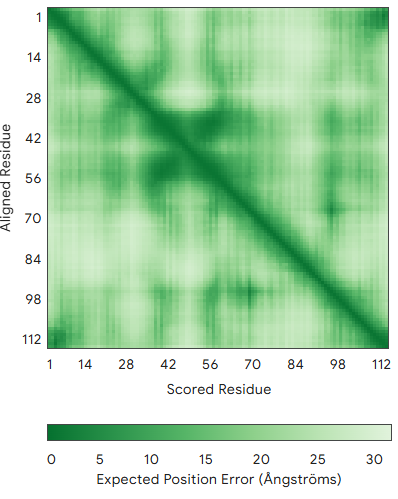

Mycgr3111636

Mycgr3111223

Mycgr3111221

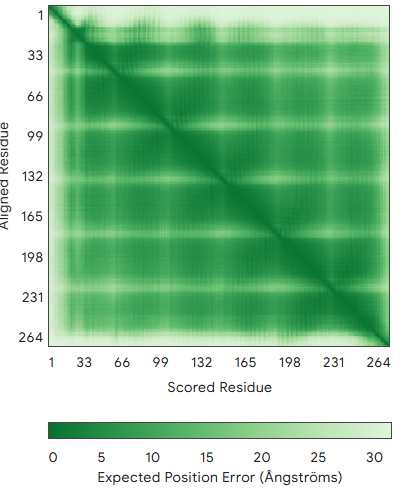

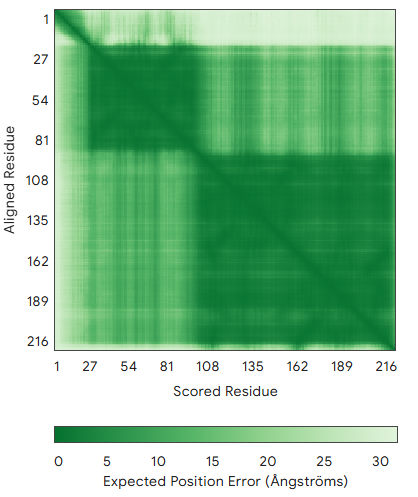

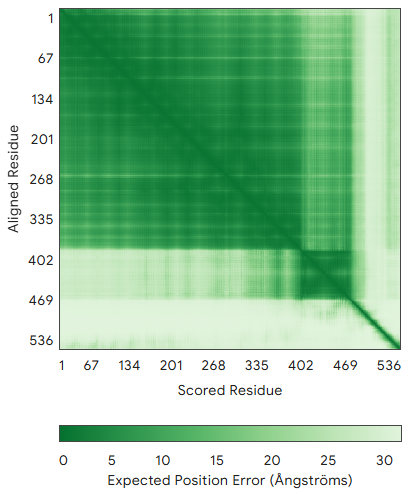

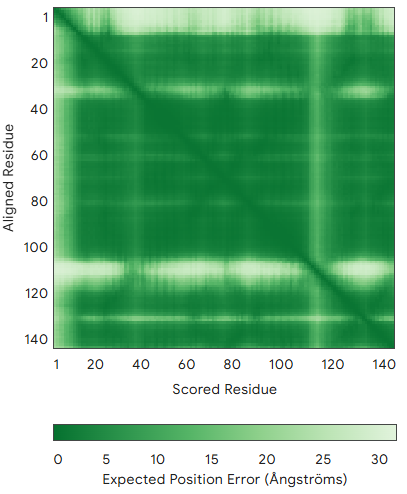

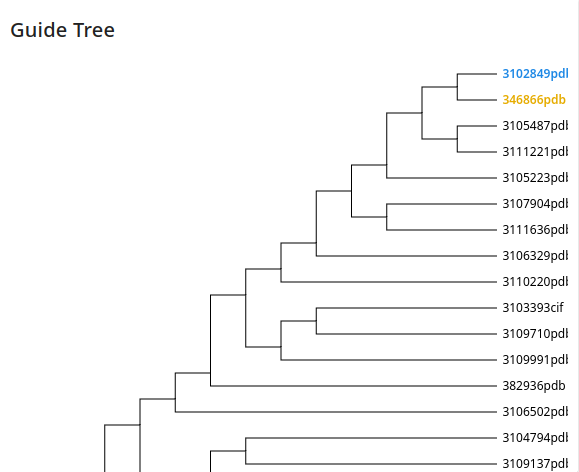

**B)**

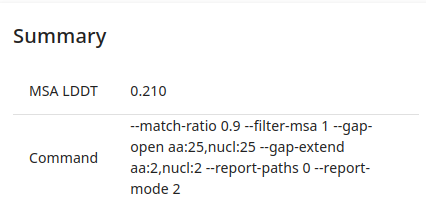

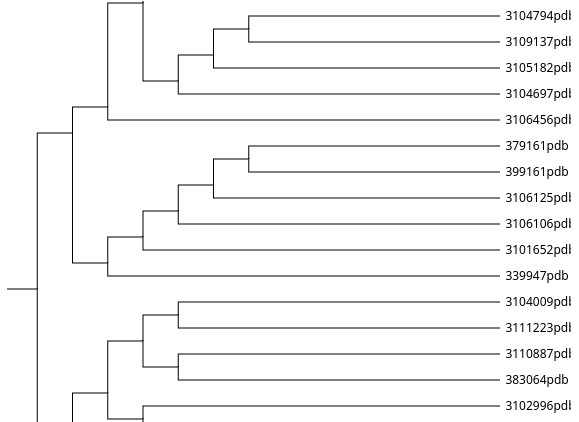

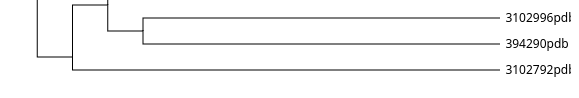

**Supplementary Figure 1.** Structural analysis of predicted effector proteins. **A.** PAE plots generated by Alphafold. **B.** Structural alignment of effector proteins using FoldMason. Effectors were grouped based on multiple structural alignment performed with FoldMason MSA (<https://search.foldseek.com/foldmason>).
