## Supplementary figure 2 for "Assessing *Zymoseptoria tritici* Candidate Effectors in the Non-Host *Nicotiana benthamiana* Reveals Immune Recognition and Potential Host-Independent Activities"

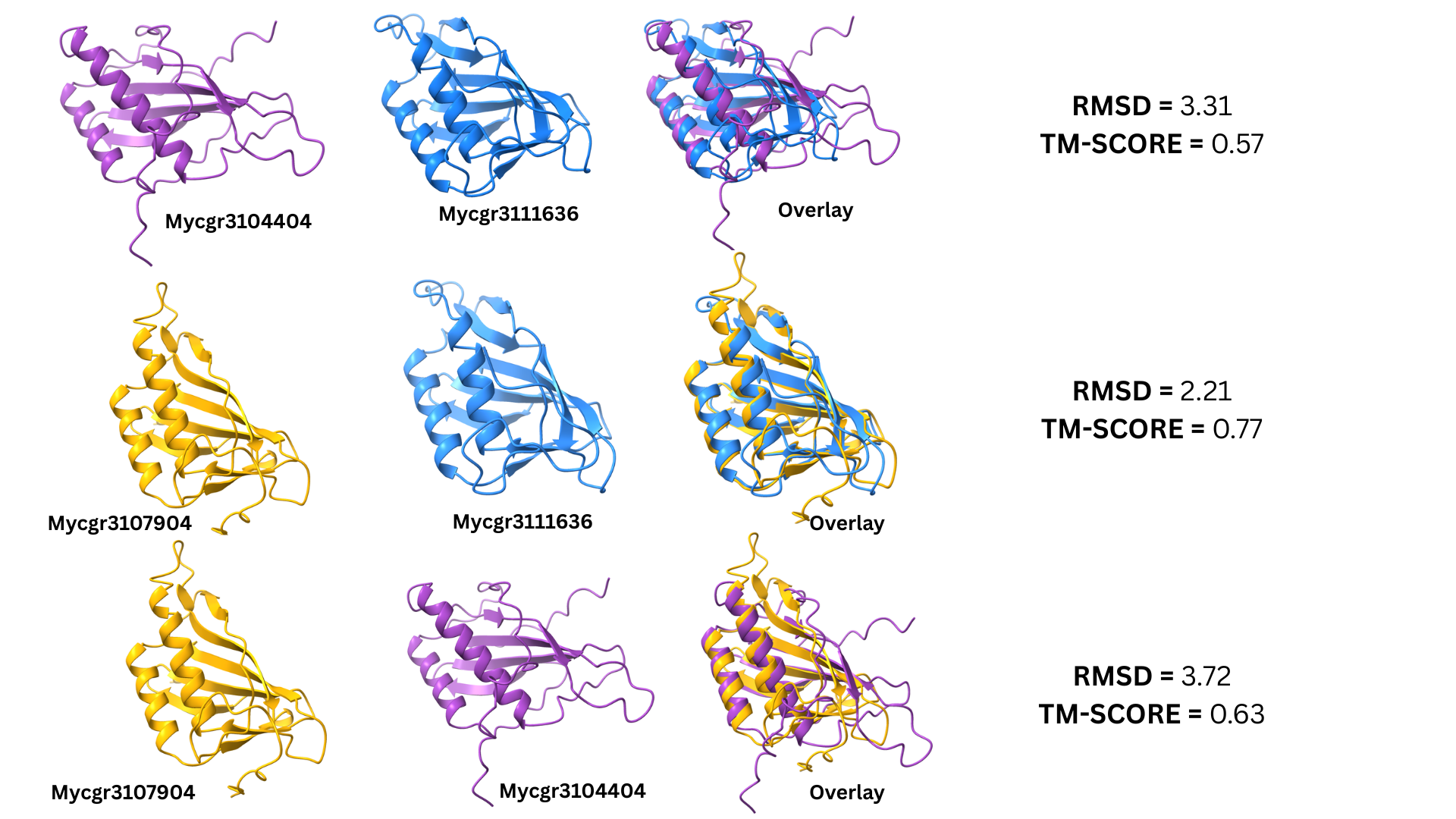

**Supplementary Figure 2.** Structural alignments of the Hce domain-containing effectors Mycgr3107904, Mycgr3111636 and Mycgr3104404.
