## Supplementary table 1 for "Assessing *Zymoseptoria tritici* Candidate Effectors in the Non-Host *Nicotiana benthamiana* Reveals Immune Recognition and Potential Host-Independent Activities"

Supplementary Table 1. Normalized count values for thirty-two candidate effector genes from *Zymoseptoria tritici* that are highly expressed in either the asymptomatic stage of infection or during the transition to the necrotrophic stage, with the highest count in bold. DAI refers to days after inoculation of *Z. tritici.*

| **ID** | **DAI 1** | **DAI 3** | **DAI 6** | **DAI 10** |
| --- | --- | --- | --- | --- |
| Mycgr3109991 | **179** | 141 | 65 | 16 |
| Mycgr3105487 | 125 | 21 | 201 | **942** |
| Mycgr3106125 | 96 | 51 | 37 | **110** |
| Mycgr3111223 | 93 | 58 | 115 | **183** |
| Mycgr31102020 | 74 | 22 | 117 | **409** |
| Mycgr346866 | 405 | **962** | 865 | 116 |
| Mycgr383064 | 343 | 34 | 144 | **977** |
| Mycgr3102792 | 345 | 210 | 2587 | **12817** |
| Mycgr379161 | **625** | 94 | 71 | 167 |
| Mycgr3106106 | **1383** | 9 | 91 | 174 |
| Mycgr3111636 | 553 | 88 | 878 | **3054** |
| Mycgr3109137 | 151 | 96 | 122 | **529** |
| Mycgr3102996 | 145 | **938** | 339 | 25 |
| Mycgr3110887 | **837** | 828 | 492 | 228 |
| Mycgr3104009 | 399 | 510 | **567** | 113 |
| Mycgr339947 | **279** | 110 | 121 | 157 |
| Mycgr3105223 | 168 | 78 | 153 | **628** |
| Mycgr3105182 | 771 | 208 | 439 | **830** |
| Mycgr3104697 | 243 | 33 | 238 | **846** |
| Mycgr3101652 | 72 | 8 | 39 | **80** |
| Mycgr3106456 | **260** | 191 | 185 | 118 |
| Mycgr3106329 | **985** | 138 | 258 | 172 |
| Mycgr382936 | 125 | 310 | 204 | **534** |
| Mycgr399161 | 260 | **404** | 268 | 339 |
| Mycgr3102849 | **1320** | 851 | 2106 | 941 |
| Mycgr3109710 | 116 | **521** | 261 | 31 |
| Mycgr3103393 | 1626 | **1631** | 1609 | 1018 |
| Mycgr3107904 | 46 | 16 | 85 | **226** |
| Mycgr394290 | **364** | 199 | 121 | 119 |
| Mycgr3106502 | **219** | 39 | 44 | 56 |
| Mycgr3104794 | 197 | 39 | 149 | **720** |
| Mycgr3111221 | 239 | 45 | 338 | **1353** |
