## Supplementary table 2 for "Assessing *Zymoseptoria tritici* Candidate Effectors in the Non-Host *Nicotiana benthamiana* Reveals Immune Recognition and Potential Host-Independent Activities"

**Supplementary Table 2.** Structural prediction and homology search results for candidate effectors of *Zymoseptoria tritici*. AlphaFold3-predicted structures were generated for 32 candidate effectors and evaluated using the predicted Template Modeling score (pTM) as a measure of overall model confidence, where values closer to 1.0 indicate higher structural reliability. Structural similarity searches were performed using FoldSeek and the Distance-matrix ALIgnment (DALI) server to identify proteins with similar three-dimensional folds in publicly available structure databases. For each candidate effector, the table reports the best FoldSeek match with its corresponding E-value (where lower values indicate stronger structural similarity), and the best DALI match with its corresponding Z-score (where values above 2 are generally considered significant). Candidate effectors with no significant structural homologs are indicated as "No match".

| **Effector ID** | **AlphaFold pTM** | **FoldSeek Best Match** | **FoldSeek E-value** | **DALI Best Match** | **DALI Z-score** |
| --- | --- | --- | --- | --- | --- |
| **Mycgr3103393** | 0.67 | SCP (Uniprot ID: N1PYP4 ) | 8.6 × 10^-25^ | SCP |  |
| **Mycgr3102849** | 0.55 | MoHrip2 (PBD ID : 5FID) | 6.2 x 10^-7^ | Elicitor Protein HRIP2 | 16.1 |
| **Mycgr399161** | 0.28 | Uncharacterized protein | 4.1 x 10^-11^ | No match | NA |
| **Mycgr382936** | 0.89 | Blue (type 1) copper domain-containing protein (Uniprot: A0A2D3UUH8) | 2.30 x 10^-20^ | Plastocyanin  (PDB ID: 1PCS) | 11.2 |
| **Mycgr3106329** | 0.12 | No structural similarity found | NA | No match | NA |
| **Mycgr3106456** | 0.80 | Yeast cell wall synthesis (Uniprot ID: M2YIF9**)** | 1.22e-^27^ | PEPTIDASE C39-LIKE DOMAIN-CONTAINING PROTEIN | 7.0 |
| **Mycgr3101652** | 0.27 | CBM1 domain-containing protein (Uniprot ID: A0A1Y6LXJ7) | 3.93e-^2^ | Antifungal protein NFAP2  (PDB ID: 8RP9) | 2.5 |
| **Mycgr3104697** | 0.80 | Small secreted protein (Uniprot ID: A0A0F4GE44) | 3.98e-^20^ | Root induced secreted protein TSP1 (PDB ID: 7CWP) | 17.8 |
| **Mycgr3105223** | 0.89 | Cell wall protein PhiA (Uniprot ID: A0A139GZZ9) | 1.49e-^15^ | No match* | NA |
| **Mycgr3105182** | 0.87 | AA1-like domain-containing protein (Uniprot ID: A0A6A6FTY4) | 4.38e-^13^ | *Alternaria* *alternata* allergen Alt A 1 (PBD ID: 3V0R) | 8.0 |
| **Mycgr3104009** | 0.73 | FAS1 domain-containing protein (Uniprot ID : A0A1V8SD58) | 2.94e-^62^ | FOURTH FAS1 DOMAIN STRUCTURE of human Bigh3 | 19.8 |
| **Mycgr339947** | 0.90 | Cerato-Platanin  (Uniprot ID M2NHG5**)** | 7.37 x 10^-20^ | MPCP2 Cerato-Platanin (3SUK) | 22 |
| **Mycgr3110887** | 0.65 | Secreted protein (Uniprot ID: A0A0F4GTW8) | 1.71e-^13^ | SUCCINATE DEHYDROGENASE [UBIQUINONE] FLAVOPROTEIN  (PDB ID : 9KPS) | 6.4 |
| **Mycgr3102996** | 0.43 | Hydrophobin (Uniprot ID: A0A1Y6LBL4) | 1.42e-^13^ | No match | NA |
| **Mycgr3110220** | 0.29 | Uncharacterized protein | 4.04e+^0^ | No match | NA |
| **Mycgr3109137** | 0.82 | Uncharacterized protein (*Fulvia fulva*) (Uniprot ID: A0A9Q8L6L7) | 9.14e-^10^ | Root induced secreted protein TSP1 (PDB ID: 7CWP) | 10.3 |
| **Mycgr3111223** | 0.71 | DnaJ-domain-containing protein (Uniprot ID: **A0A316US27)** | 3.56e-^23^ | DNAJ homolog subfamily C member 3  (PDB ID: 3IEG) | 30.4 |
| **Mycgr3111636** | 0.79 | Ecp2 effector protein-like domain-containing protein (Uniprot ID: A0A9P4I4X9) | 1.30e-^7^ | HCE2 domain-containing protein (PDB ID: 8ACX) | 16.6 |
| **Mycgr3106106** | 0.89 | Uncharacterized protein (Uniprot ID: A0A2P7ZE25) | 6.24e-^13^ | MYCOBACTERIUM SMEGMATIS MFD  (PDB ID: 6ACX) | 4.5 |
| **Mycgr346866** | 0.58 | Lytic polysaccharide monooxygenase (Uniprot ID: A0A4U0X666) | 1.20e-^46^ | AA11 FAMILY LYTIC POLYSACCHARIDE MONOOXYGENASE B  (PDB ID: 9HDG) | 30.5 |
| **Mycgr379161** | 0.35 | Chitin-binding type-2 domain-containing protein (Uniprot ID: **A0A9D4LDY0)** | 7.93e+^0^ | No match | NA |
| **Mycgr3102792** | 0.49 | Uncharacterized protein (Uniprot ID: A0AAD8YHW2) | 5.26e+^0^ | DNA-DIRECTED RNA POLYMERASE I SUBUNIT RPA190 | 5.0 |
| **Mycgr383064** | 0.57 | CBM1 domain-containing protein (Uniprot ID: A0A1X7S9K2) | 4.31e-^5^ | No match | NA |
| **Mycgr3107904** | 0.79 | Ecp2 effector protein-like domain-containing protein (Uniprot ID: G9NEU2) | 2.28e-^9^ | HCE2 DOMAIN-CONTAINING PROTEIN | 18.4 |
| **Mycgr3106125** | 0.62 | Hydrophobin (Uniprot ID: A0A2H1H3R4) | 4.32e-^5^ | MU-DIGUETOXIN-DC1A | 2.6 |
| **Mycgr3109710** | 0.93 | Pry1 CAP domain  (PDB ID: 5JYS) | 1.81e-^11^ | PROTEIN PRY1  (PDB ID : 5JYS) | 18.8 |
| **Mycgr3105487** | 0.81 | Cladosporium fulvum LysM effector Ecp6  (PDB ID: 4B8V) | 1.46e-^4^ | CLADOSPORIUM FULVUM LYSM EFFECTOR ECP6  (PDB ID: 4B8V) | 8.6 |
| **Mycgr3109991** | 0.13 | No match | NA | No match | NA |
| **Mycgr3104794** | 0.82 | AA1-like domain-containing protein (Uniprot ID: A0A6A6CDX9) | 4.10e-^9^ | ROOT INDUCED EFFECTOR PROTEIN TSP1 | 9.7 |
| **Mycgr394290** | 0.14 | 60S ribosomal export protein NMD3 (Uniprot ID: A0A7S1F7B3) | 8.50e-^2^ | No match | NA |
| **Mycgr3106502** | 0.23 | Long chronological lifespan protein 2 (Uniprot ID: A0A2H1H8A4) | 4.86e-^8^ | No match | NA |
| **Mycgr3111221** | 0.64 | Cladosporium fulvum LysM effector Ecp6  (PDB ID: 4B8V) | 3.13e-^28^ | CLADOSPORIUM FULVUM LYSM EFFECTOR ECP6  (PDB ID: 4B8V) | 24.5 |
